# Divergence of an extracellular contractile injection system infectivity elucidated by high resolution structural studies of its tail-baseplate complex

**DOI:** 10.1101/2025.07.21.665459

**Authors:** Anindito Sen, Tsukasa Nakamura, Genki Tarashi, Veer Bhatt, Kyungho Kim, Shankararaman Chellam, Le Tran, Daisuke Kihara

## Abstract

Divergence to the infectivity practice by extracellular contractile injection systems (eCISs) is displayed by myophage P1 in its lytic phase involving its baseplate receding away from the host bacterium during infection. Atomic structure of the proteins forming P1’s Tail Baseplate (TB) complex, determined here, using cryo electron microscopy are employed to identify a sequence of viral events explaining this unique phenomenon. The P1 baseplate is found to be devoid of protein appendages that are necessary for anchoring it to the host bacterium. To compensate this deficiency, P1’s Long Tail Fibers (LTFs) affix to the host’s exterior, straighten up imparting stability to the virion for pursuing the infection process, thereby eliciting the baseplate hub and the tail sheath to ascend away, resulting in uniform compression of the latter. Upon the maximum unkinking of LTFs, a descending corkscrew motion commences that results in the non-uniform downward compression of the tail sheath, the tail tube and baseplate needle that perforates through the host’s membrane-cytoplasm, leading to a successful phage-bacterium infection.

## Introduction

Extracellular contractile injection systems (eCISs) are nano-injections that deliver payloads to sub-cellular targets by pursuing a well-studied infectivity procedure (1–14). All eCISs like P1 myophage TB complex is an amalgamation of a highly contractile sheath encapsulating a rigid non-contractile tail tube that terminates on the top of a 6-fold symmetrical, structurally complex baseplate that possesses six LTFs (1) (Fig. S1A). During phage-bacterium infection, a myophage particle, attaches to the outer surface of the host cell using LTFs, then firmly anchors its baseplate by protein appendages called **s**hort **t**ail **f**ibers (STFs) resulting in significant conformational changes in the baseplate (2). The tail sheath undergoes a non-uniform contraction, triggered by the conformational changes in the baseplate, propelling the tail tube downwards in a corkscrew manner, along with the capsid. A needle, positioned at the distal end of the tail tube punctures through the cell membrane, disassociates itself, leading the tail tube into the host cell cytoplasm releasing the phage genome inside for replication (3–14). This entire process of the infection has been studied extensively for T4 myophage and is widely regarded as the sole infection mechanism for eCISs (and denoted as CIS from here on). However, cryo-electron tomography (cryo-ET) studies of the P1 infection process indicated a divergence of this mechanism (1). The results showed a noteworthy movement of the hub of the P1 baseplate away from the outer membrane of the host cell during the infection process, which is unheard of in other CISs. To investigate this unique viral phenomenon, we have built de novo atomic structures for 12 unique gene products of P1 tail proteins (except for the LTFs) in its pre- and post-contracted states by employing cryo-Transmission Electron Microscopy (TEM) (Fig. 1, SI 1). We studied the interactions of the different protein subunits of the TB complex and employed their atomic structures to build a sequence of viral events to derive an explanation comprising a dual act of the LTFs and tail sheath. The modus operandi of the P1 TB complex explained here successfully delineates its unique infection mechanism at the atomic level and establishes that a minimalistic contact between the virion and its host solitarily through the LTFs is good enough to decimate the later, making it a perfect bio nano carrier of therapeutics to be delivered to desired sub-cellular targets with high precision.

**Fig. 1.**
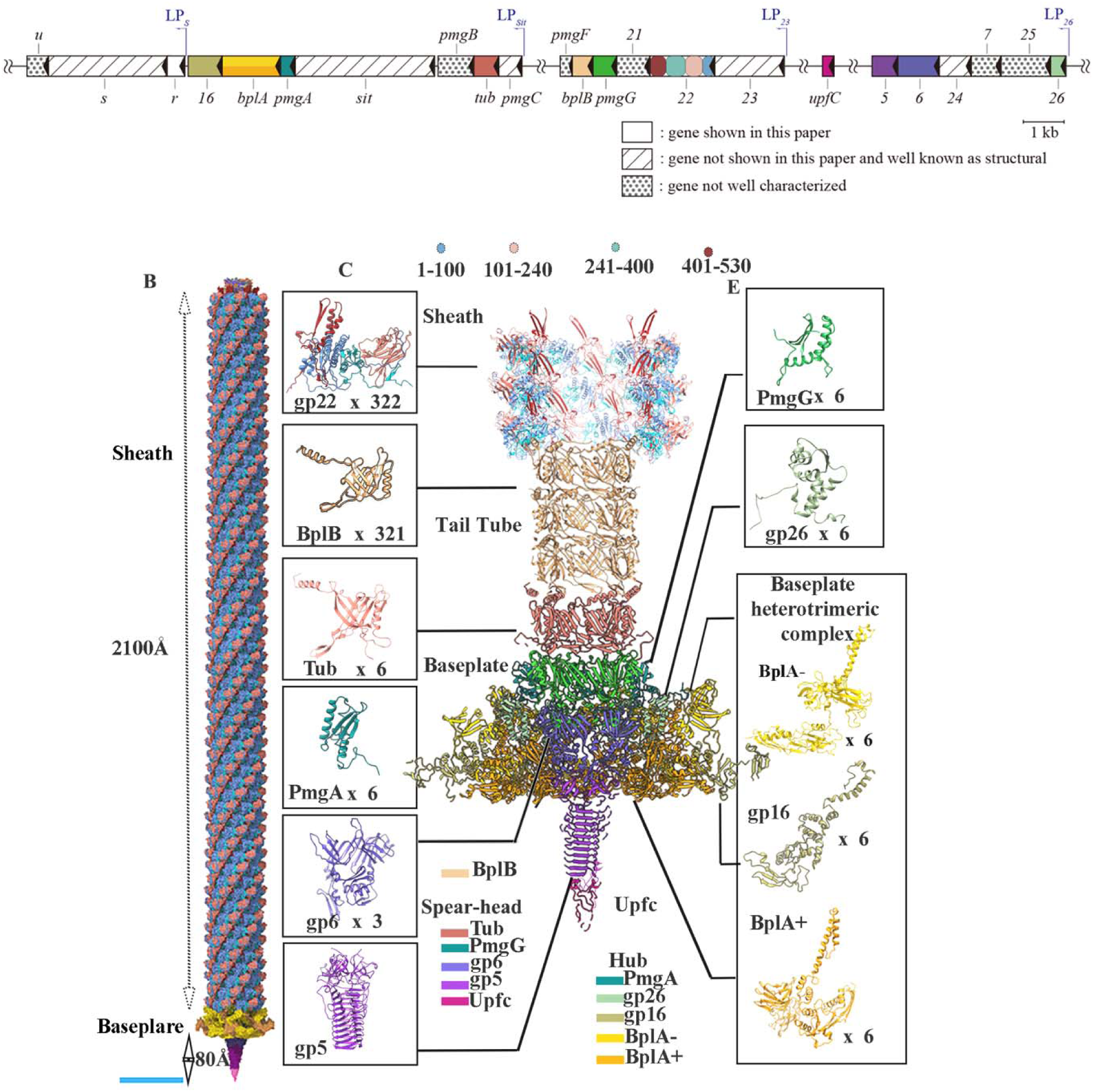
Gene diagram and atomic structure of the Pl phage TB proteins. **(A)** Organization of the TB protein genes with their neighboring genes. Boxes with internal triangles show the positions on the P1 genome and orientations of the genes with their names. Colors of the boxes correspond to the colors of the structures shown in (C). Gene boxes without hatch are genes shown in this paper. Boxes with slashes are genes not shown in this paper and are well known as structural proteins in previous studies (22). Boxes with dots are genes that are not well characterized. Thin deep blue half arrows indicate the start sites and directions of late promoters. **(B)** Iso-surface representation of the entire TB complex without the LTFs. The sheath shields nearly 96% of the entire TB complex. Bar: 200Å. (**C**) shows the atomic structure TB complex along with the single subunit of each of the proteins. The total number of copies of each of the protein subunits is written as the ‘x copy number’. Each of the proteins, except gp22 and the tail tube protein (BplB), is categorized in two groups: the spearhead and the hub regions and is given a different color (shown below the baseplate). The gp22 is divided into 4 different segments each color coded for a range of residues mentioned on the top of (C). The categorization is done based on occurrence and their roles in the infectious process. The residues of gp22 (1–100) formed the inner region of the sheath tube with negative and neutral electrostatic charges that help the tail tube to get pushed down during the infection process. Residues 100-240 for the outer globular domain of the sheath and are electronegative in nature, while 241-400 forms the middle part of the sheath and holds the fourth segment, residues 401-529, which forms the central section of the sheath tube. This segment electrostatically holds the sheath molecules to the tail tube and participates in the handshake domain holding the gp22 subunits together even during the contraction. The ten baseplate proteins are Tub, PmgA, gp6, gp6, PmgG, gp26 along with the triplex BplA+, BplA- and gp16.

## Structure of P1 TB complex

### The contractile sheath

One of the principal sections of the P1 TB complex is its 18.3 megaDalton contractile helical hollow tube 2100 Å long tubular sheath tube, formed by 322 copies of protein gp22, following a strict helical rule throughout 96% of the entire sheath tube length (SI T2). The last two slabs are however slightly skewed when compared to the rest of the hexagonal sheath slabs thus violating the helical rule of the pre-contracted sheath tube, (Fig.S1) (4). The gp22 molecule is divided into 4 segments with the residues 1-100 forming part of the inner region (domain 1a), 101-240 forming the highly electronegative globular domain 2 followed by domain 1b with residues 241-400 forming part of the internal and external central domain that acts as an amalgamation between the globular and the 3^rd^ domain (residues 401-529) (Movie S1). This 3^rd^ domain participates in the weak interaction between the sheath and tail tube. Domains 1,3 are mixed combinations of positive, negative and neutral charged residues with each of them playing significant roles in the architecture of the TB complex and infection process. Two consecutive coplanar sheath subunits with residues 15-18 and residues 520-523 forming -strands oriented in parallel, interact with the two anti-parallel -strands (residues 494-502 and residues 505-512) of sheath subunit belonging to the lower slab creating the handshake motif that is responsible for a unique 3-way interconnectivity among the subunits which is always preserved during the sheath contraction. Unrolled sections of the density maps reveal the significant reorganization of the sheath molecules as the inclination of the protofilaments changes by >20° with the horizontal yet maintaining the 6-fold point group symmetry among the gp22 molecules (Fig. 2A, Fig. S2A-B). Computational morphing among the pre- and post-contracted states of gp22 discloses that the gp22 subunit rotates and extends outward as a single entity except residues 1-35 and 520-529, which twist and displace like an arm to gyrate the rest of the gp22 molecule. This act maintains the handshake domain during the contraction and the sheath molecule integrity even during massive conformational changes (Fig. 2B, Movie S2). The conformational changes in the sheath subunits due to the contraction, result in the increase of electronegative potential inward, directed towards the tail tube, suggests that the sheath contraction debilitates the electrostatic forces holding the inner tube to the sheath tube (Fig. S2C).

**Fig. 2.**
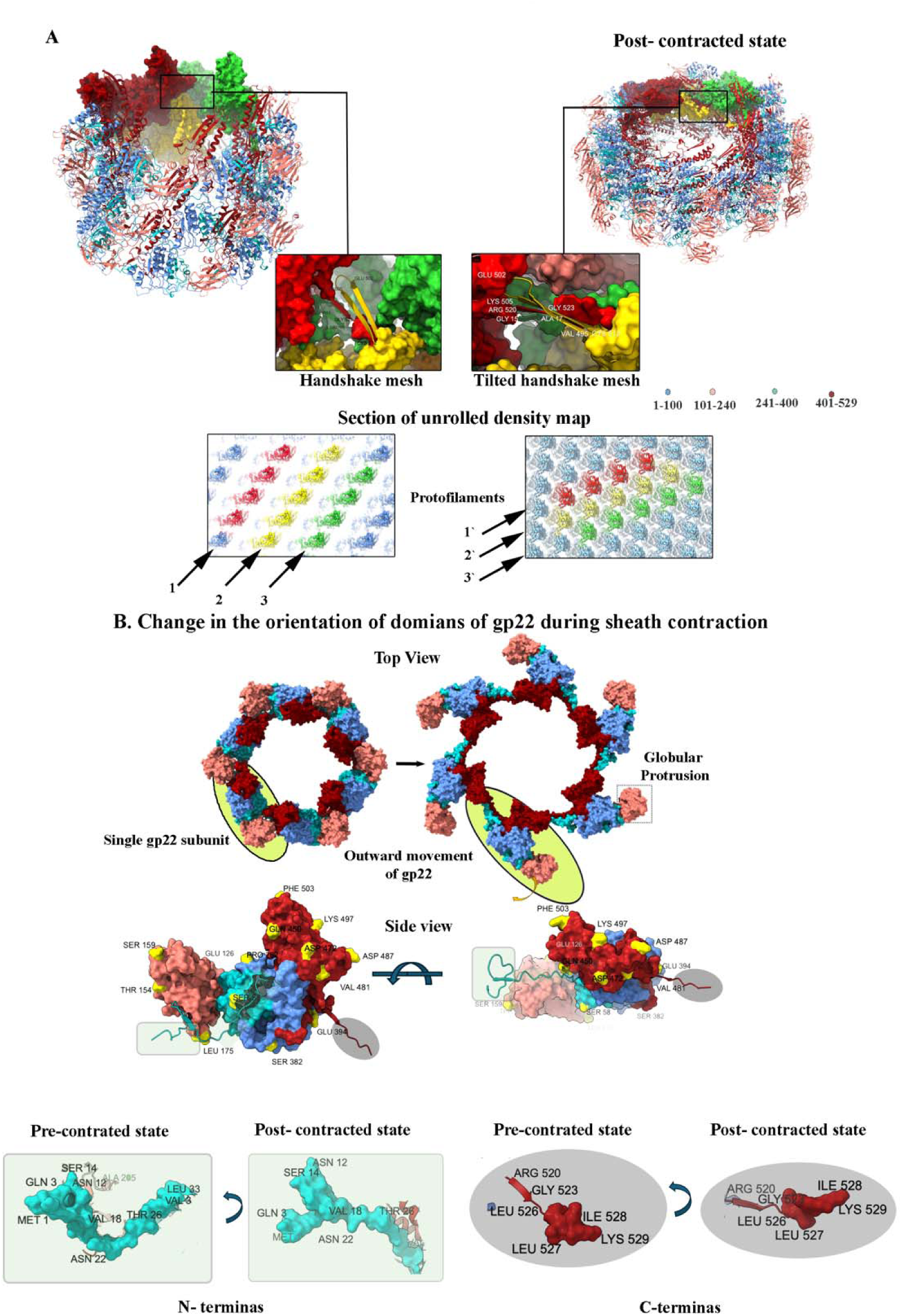
Atomic structure of sheath tube of Pl phage in its pre- and post-contracted states. **(A)** Atomic structure of normal and compressed states of tail sheath (shown in an inclined manner w.r.t to the vertical helical axis). Three subunits are colored in red, yellow and green and the corresponding handshake domains in the pre- and post-compressed states are shown below. Residues 432-477 are removed from the side views of the two states and the handshake motifs are encircled for better visualization. The ß-strands of the handshake motif tilt as the sheath compresses but preserve their alliance. Sections of the electron density maps are computationally unrolled to show the arrangement of gp22 molecules (the globular domains). Three protofilaments colored red (1), yellow (2) and green (3) show the change in their orientation going from a normal to a compressed state. The color codes show the different domains of gp22. **(B)** Change in the orientations of the domains due to sheath contraction. Top views of the atomic structures of a single ring of the sheath tube in its pre- and post-contracted states showing the sidewise expansion of the sheath tube with one gp22 molecule highlighted in either of the states. A gp22 molecule, with several residues colored in yellow and labelled to show gyration of the different domains. The arcs arrows show the direction of the gyration. The main body of gp22 (residues 36-520) rotates as a whole, without independent flexing of any of its domains except the N-terminus (residues 1-35, colored with light green background), and the end region of the C-terminals (residues 521-529, shown with dark grey background) twists, but significantly more that main body (Movie S2), thus maintaining the handshake mesh, upholding the interactions among the subunits transitioning to a newer helical organization, upholding the sixfold point-group symmetry at all times of the compression.

### The P1 tail tube

The P1 helical tail tube is 5.7 megadalton protein complex and a gene product of BplB which has cross -strands motif at the center with two extended regions on one side, following helical symmetry identical to the sheath tube (Fig. 3A, S3A, SI T2). The subunits forming the tube helix are electrostatically attached to the neighboring subunits by a unique 4-point interaction. The extended N-terminal region of BplB subunit is a combination of positive and neutral electrostatic charges which travels between 2 perpendicular tail tube slabs of highly electronegative charged regions, forming strong electrostatic bonds. These ‘arms’ of the tail subunit act like a ‘sealant’ holding its neighboring subunits of the upper and lower tiers together (Movie S3). As shown in Fig. 3B, residues Lys11 and Lys118 of the subunit ‘e’, colored in violet, interact with Asp169 of subunit ‘a’ and Glu90 of subunit ‘d’ respectively. Conversely, residues Asp169 and Glu90 of subunit ‘e’ interact with Lys11 and Lys 118 of subunits ‘i’ and ‘f’, respectively. These strong inter-subunit electrostatic interactions contribute to the non-flexibility of the tail tube. Investigation of the interaction of the tail tube and sheath subunits reveals no physical contacts between them, but electrostatic interactions hold them in position (Fig. 3C, Fig. S3B). The negatively charged region of the tail tube subunits Glu133 and Glu167 interact with Lys465 and Lys469 of gp22, respectively. The electrostatic interaction among the tail tube and sheath subunits is not overwhelmingly strong throughout the sheath-tail tube interface but is restricted to ‘pockets’ of regions. The outer region of the tail tube is predominantly negatively charged with bands of positive charges at the end of each hexagonal slab. The internal region of the P1 tail tube is a combination of electronegative and neutral charge distributed in a spiral manner, suggestive of the fact that the inner layer of the tail tube acts as a lubricant for the phage genome to travel down to the tip of the tail tube to be released in the host cytoplasm for replication (Fig. S3C).

**Fig. 3.**
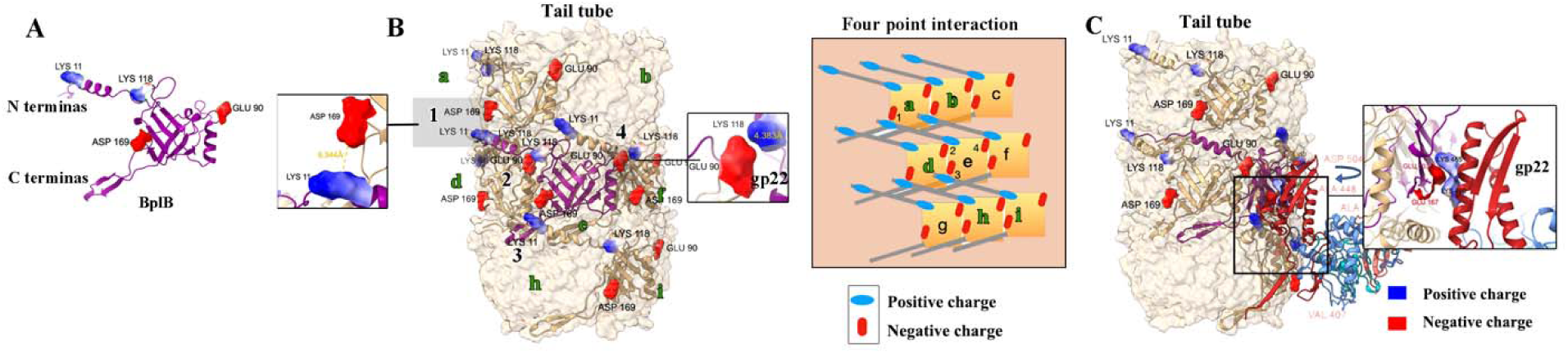
Tail tube and sheath subunits electrostatic interaction. **(A)** Atomic structure of BplB subunit with surface representation of the residues, responsible for the interaction with their neighboring subunits, along with their charges are shown. **(B)** 4-point electrostatic interactions of the BplB subunit, colored in violet, labelled ‘e’, among its neighboring subunits, labelled ‘a to i’. Distance between residues participating in the subunit interactions: Asp169 to Lys11 of the neighboring subunits is approximately 9.3Å while Lys118 to Glu90 is 4.3Å. A simplified schematic diagram with the interaction positions numbered 1-4 for the subunit ‘e’ is shown to explain the communication among the subunits**. (C)** Electrostatic interaction of BplB with gp22 and close view of the interaction of tail tube subunits Glu133, Glu167 interact with Lys465, Lys469 of sheath tube.

### The P1 baseplate

A total of 10 proteins build up the major portion of the baseplate, which can be divided into two parts, termed here as ‘spearhead’ and ‘hub’ segments (Fig. 4A-B, Fig. S4A, Movie S4). The spearhead section initiates beneath the rearmost BplB slab where six copies of Tub protein are arranged in a 6-fold symmetrical manner and concentric to the tail tube. Beneath the Tub protein, resides another hexameric slab formed by PmgG protein, a homolog to the gp48 protein of T4 bacteriophage. It has a reverse sub-conical structure whose orifice constricts from 90 Å to 60 Å. This is a unique feature as the next in-line protein complex is a trimer of gp6 whose apex is 70 Å wide and the tapered segment of PmgG fits well in it. The gp6 trimer narrows down to 50 Å at its distal end where the baseplate plate needle originates. All the 4 proteins, BplB, Tub, PmgG, and gp6 have cross -strands motif as a common structural feature irrespective of their symmetries and very different nucleotide sequences suggesting that the polymerization of the tail tube occurs in the pipeline of the formation of tail tube. The subunits of BplB, Tub, PmgG, gp6 and gp5 proteins are held to each other by electrostatic forces (Fig. 4B). However, interaction between PgmG - gp6 is not very clear from the current study owing to the paucity of much higher resolution of the baseplate density map. The needle, a product of gp5, is a reverse trimeric conical shaped assembly formed by 11 horizontal layers of -strands with each layer having three strands at a given plane. The tip of the needle is formed by UpfC protein that has six - strands and the rest as unstructured loops (Fig. 1C).

**Fig. 4.**
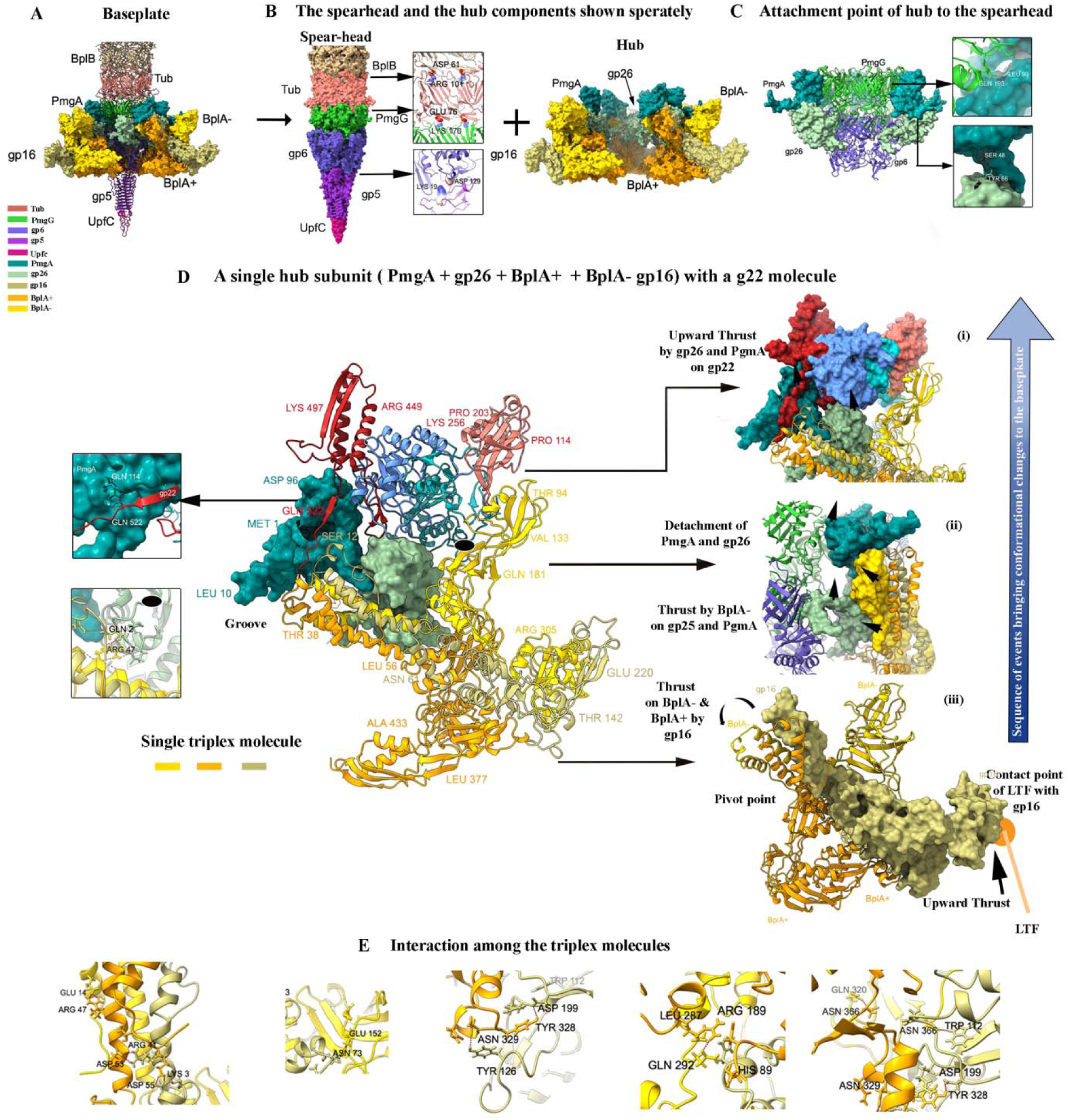
Organization of the different proteins forming the Pl baseplate. **(A)** Atomic structure of the baseplate with a copy of subunits of PmgA, gp26, gp16, BplA+ and BplA-removed for better visualization. The color codes for the different proteins involved are also shown on the left side of the image. **(B)** The baseplate is divided into sections - Spearhead and the Hub. The bottom view of the Hub region shows a three-fold symmetry among gp16, BplA+ and BplA-molecules. A single triplex unit, shown in lighter density in the baseplat bottom view, is computationally separated outward to show the orientation of gp16, BplA+ and BplA-w.r.t each other. Proteins of the spearhead regions are primarily held to each other by electrostatic forces shown for BplB, Tub, PmgG and gp6 and gp5 with the participating residues colored according to their charges, on the left. **(C**) The hub is held to the spearhead by 6 copies of PmgA and gp26 each, bound to PmgG and gp6, respectively. **(D)** A single subunit of the Hub region with a single gp22 subunit of the last sheath ring is shown. Interactions among PmgA with gp22 and BplA-with gp26, are shown separately on the left. Enlarged views show the arrangement of BplA+, gp16, BplA-in a sequential manner with gp26, PmgA and gp22 are shown on the right-hand side. Starting from the bottom, gp16 suffers initial upward thrust from the straightening of the LTFs (iii) and imparts it to BplA-making PgmA and gp26 detach from the attachment of PgmG and gp6 respectively (ii). The subunit of PgmA and gp26 is strategically positioned in close vicinity of domain 1a of gp22 (residues 1-100). The axial thrust on PgmA and gp26 is imparted to gp22 leading the sheath to compress (i). The black arrow heads show the direction of the trust and the black arc arrows show the direction of movement of the protein subunits. The distance between BplA- and gp26 is around 10Å. **(E)** The different H bonds holding gp16, BplA+ and BplA-to each other (Movie S5).

Induction of the hub components of the baseplate starts with PmgA protein, an ortholog of gp25 of the T4 phage, surrounding the PmgG interacting using weak Hydrogen (H) between Leu90 and Gln193 respectively (Fig. 4C). On the outer side, Gln114 of PmgA hold H-bonds with Gln522 of the last ring of gp22 subunit and act somewhat like a frail holder of lower portion sheath to the spearhead section. At the proximal end sits a hexameric set of gp26 molecules, each strongly held to six PmgA molecules by the interaction between Try66 (gp26) and Ser48 (PmgA) as shown in Fig. 4C. The outer region of gp26 embodies 3 proteins gp16, BplA-and BplA+, the latter two, are homolog of inner baseplate protein gp6 of T4 phage. These three protein molecules are organized to form a trimeric module, a structural pattern highly conserved among CISs (Fig. S4) (Movie S5) (2, 4–6, 11). A combination of a single of the trimeric module along with a copy of PmgA and gp26 each form a single subunit of the baseplate Hub (Fig. 4D). The N-terminal domains of the trimeric molecules, named ‘core bundle’, recline in a groove configured by residues 26-35 of PmgA. The overall architecture looks like a ‘ball-and-socket joint’ allotting significant expanse to maneuver for the trimeric module and is extremely vital for the infection process, discussed later (Fig. 4D). Each of these 3 molecules are held to each other by strong salt bridges among them with BplA+ acting like a ‘linchpin’ at the core, BplA-forms the outer rim of the baseplate while gp16 emerges as a ‘lever’ (Fig. S4B) The central region of these molecules act like a ‘pivot point’ for their conformational movements, explained in the next section (Fig. 4iii). The residues Lys3, Arg47 of BplA+ interact with Asp53 (gp16) and Glu14 (BplA-) respectively. Other connections are at the central region of the module where Tyr328 (BplA+) interacts with Asp199 (gp16) making a strong bond and Arg189(BplA+) interacts with Gln292(BplA-). Interactions between residues Asn73 and Trp112 of gp16 interact with Glu152 and Asn366 of BplA-, forming a "NH-π" hydrogen bond. The trimeric module is held to the spearhead by gp26. Residue Arg47 (BplA-) makes strong salt bridge with Asp86 of gp26, Lys160 of BplA+ forms bonds with Gln117 of gp26 and Arg9 of gp16 interacts with Asp62 of gp26 (Fig. 4E). The 6 copies of gp26 molecules interact with 3 copies of gp6 and there exists a symmetry mismatch between the two and in our current study, these interactions are not very well concluded (Fig. 4C). In-depth analysis shows no significant sequence similarity between gp16 of P1 and gp7 of T4 phage was found, yet they have much structural likeness (Table 1). The electron density P1 baseplate map revealed no protein appendages like STF network (gp9-12) of T4 are present, which would be necessary for the P1 baseplate to clinch the host cell membrane during infection (2). Furthermore, our meticulous profile-profile sequence search could not find any homolog candidates corresponding to T4’s gp9, gp10, gp11, gp12 thus supporting our findings.

**Table 1.**
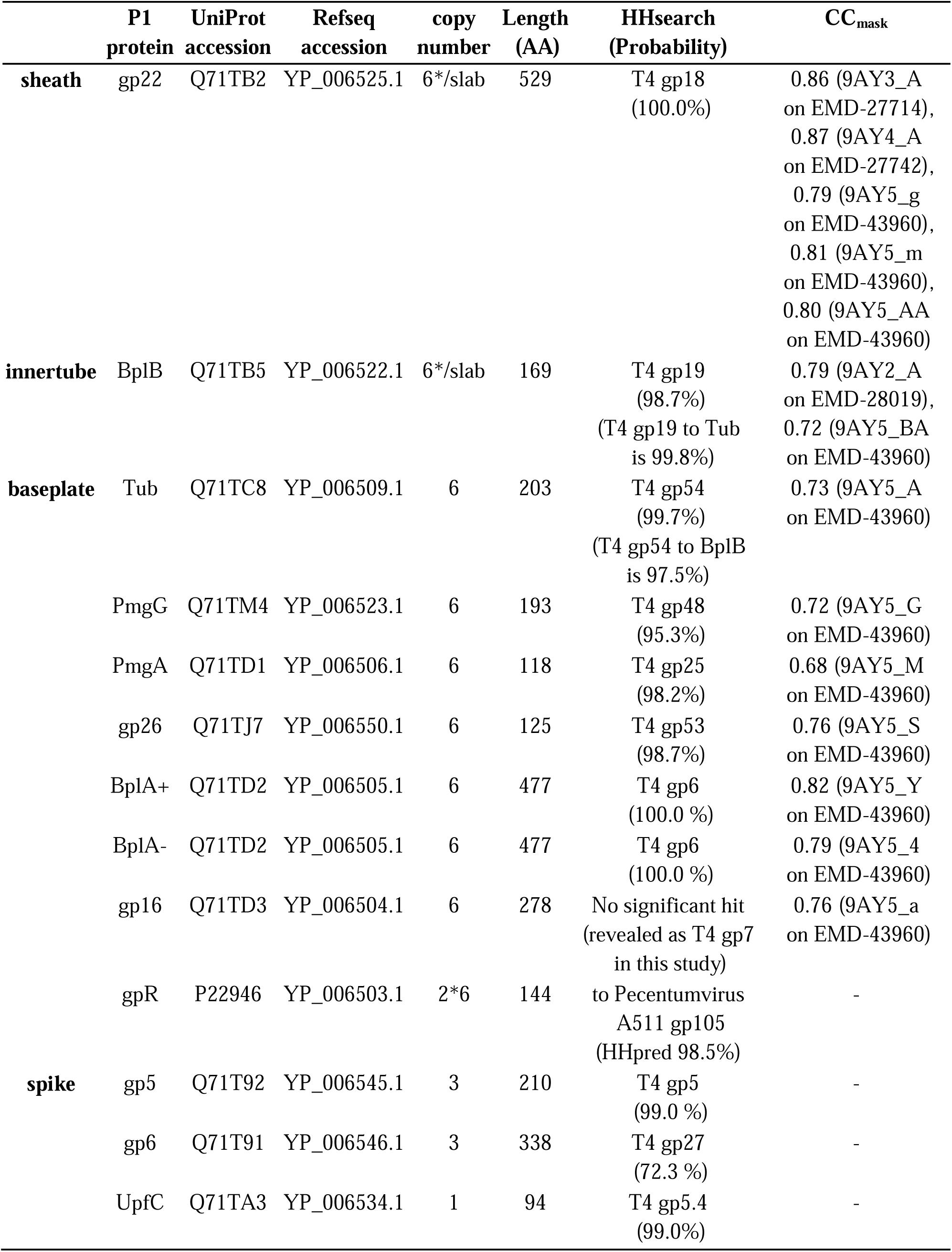
Summary of P1 phage protein sequences used for atomic structure modeling, HHsearch (19) results were calculated using hidden Markov model profiles from T4 phage sequence as a query and against all profiles from known P1 phage proteins in UniProt except duplicates and long (120 profiles total). Probability from HHsearch is a probability that a query T4 is at least partly homologous with a protein of P1 phage. About gpR, though HHsearch had no significant hit, Pecentumvirus A511 gp105 was a significant hit using HHpred (20) and it is a known ortholog of T4 gp8 (12). About gp16, HHsearch and HHpred had no significant hit but we revealed T4 gp7 is homologous in this study. The model-map correlation coefficients with the masked map around the model, CC_mask_ values, were calculated for pairs of one chain model with the earliest chain ID among chain models from each P1 protein and the corresponding volume map using Comprehensive validation tool in Phenix to confirm the qualities of modeled structures (21).

|  | P1<br>protein | UniProt<br>accession | Refseq<br>accession | copy<br>number | Length<br>(AA) | HHsearch<br>(Probability) | CC<sub>mask</sub> |
| --- | --- | --- | --- | --- | --- | --- | --- |
| **sheath** | gp22 | Q71TB2 | YP_006525.1 | 6*/slab | 529 | T4 gp18<br>(100.0%) | 0.86 (9AY3_A<br>on EMD-27714),<br>0.87 (9AY4_A<br>on EMD-27742),<br>0.79 (9AY5_g<br>on EMD-43960),<br>0.81 (9AY5_m<br>on EMD-43960),<br>0.80 (9AY5_AA<br>on EMD-43960) |
| **innertube** | BplB | Q71TB5 | YP_006522.1 | 6*/slab | 169 | T4 gp19<br>(98.7%)<br>(T4 gp19 to Tub<br>is 99.8%) | 0.79 (9AY2_A<br>on EMD-28019),<br>0.72 (9AY5_BA<br>on EMD-43960) |
| **baseplate** | Tub | Q71TC8 | YP_006509.1 | 6 | 203 | T4 gp54<br>(99.7%)<br>(T4 gp54 to BplB<br>is 97.5%) | 0.73 (9AY5_A<br>on EMD-43960) |
|  | PmgG | Q71TM4 | YP_006523.1 | 6 | 193 | T4 gp48<br>(95.3%) | 0.72 (9AY5_G<br>on EMD-43960) |
|  | PmgA | Q71TD1 | YP_006506.1 | 6 | 118 | T4 gp25<br>(98.2%) | 0.68 (9AY5_M<br>on EMD-43960) |
|  | gp26 | Q71TJ7 | YP_006550.1 | 6 | 125 | T4 gp53<br>(98.7%) | 0.76 (9AY5_S<br>on EMD-43960) |
|  | BplA+ | Q71TD2 | YP_006505.1 | 6 | 477 | T4 gp6<br>(100.0 %) | 0.82 (9AY5_Y<br>on EMD-43960) |
|  | BplA- | Q71TD2 | YP_006505.1 | 6 | 477 | T4 gp6<br>(100.0 %) | 0.79 (9AY5_4<br>on EMD-43960) |
|  | gp16 | Q71TD3 | YP_006504.1 | 6 | 278 | No significant hit<br>(revealed as T4 gp7<br>in this study) | 0.76 (9AY5_a<br>on EMD-43960) |
|  | gpR | P22946 | YP_006503.1 | 2*6 | 144 | to Pecentumvirus<br>A511 gp105<br>(HHpred 98.5%) | - |
| **spike** | gp5 | Q71T92 | YP_006545.1 | 3 | 210 | T4 gp5<br>(99.0 %) | - |
|  | gp6 | Q71T91 | YP_006546.1 | 3 | 338 | T4 gp27<br>(72.3 %) | - |
|  | UpfC | Q71TA3 | YP_006534.1 | 1 | 94 | T4 gp5.4<br>(99.0%) | - |

It is unknown exactly where the LTFs are anchored to the P1 baseplate, either directly or via some protein/s as in T4 phage (2). The baseplate density map at a very low threshold exhibits presence of some weak densities on the distal end of gp16 suggesting that they are very flexible and are averaged out of the electron density map, could potentially be the point of attachment of the LTFs to the baseplate and is supported by the Two-Dimensional class average of the images of the baseplate (not shown here) that revealed some weak fuzzy densities attached to the baseplate rim (Fig. S4C). Similar observations were made on the structure of phage E217 where the LTF is found directly attached to the triplex subunit gp45/gp44-ab alike gp16/BplA+-for P1 phage, forming the rim of baseplate (11). For T4 phage, LTFs bind to the baseplate through a trimer of gp9 that attaches to gp7, which is also associated with gp8, a part of the trimeric complex of the intermediate baseplate (2). Given the close resemblance of P1 hub proteins with the inner and parts of the intermediate baseplate of T4, and the presence of feeble densities over the gp16 in the P1 baseplate-tail tube density map, we predict with some credence that the place of contact for the LTFs to the P1 baseplate is through gp16 (2). Our search also revealed that protein gpR of P1 phage has indirect similarity to gp8 of T4 phage (Table 1). Using the sequence, a predicted structure of gpR is developed whose dimer showed good structural interaction with gp16, similar to an arrangement in T4 phage with gp8 and gp7 (Fig. S4C) (2). We anticipate that the LTFs in P1 can fasten to the dimer of gpR and/or to the unmodeled region (240–278) of gp16 directly, or through some unknown protein.

### The P1 TB dynamics triggering infection process of P1

During infection, P1 LTFs fasten themselves to specific receptors located on the external surface of the host cell membrane. This positions the virion in its pre-contracted state, 530 Å from the host’s outer surface (Fig. S4D, S5A) (1, 6). In the absence of STFs, the anchoring LTFs straighten up from their kink regions to provide adequate mechanical support for the virion particle to remain attached to the host cell body during the entire infection process and exert an upward thrust, transmitted through gpR to gp16 and subsequently to BplA- and BplA+. As the other ends of the triplet i.e. the N-terminal domains reside in the groove region, the upward motion is not contested by any other section of the baseplate (Fig. 4Diii). This upward movement causes BplA-to exert an upward mechanical thrust to gp26, being closest to it (10Å), and to PmgA, disengaging both from the spearhead therefore propelling the gp22 molecules upward causing the sheath to contract, launching prodigious conformational changes in the entire TB complex (Fig. 4D(ii-i), 5A). The conformational changes in the P1 baseplate are supported by the reconstructed density maps in the cryo-ET work, which shows an expansion of the baseplate’s width to 280 Å during infection (1). This observation suggests that the hub proteins have traversed enough distance upward to exert a significant mechanical thrust to compress the sheath tube. Similar baseplate changes are also reported for E217 phage, thus supporting our findings (11). This very detachment of the baseplate hub from the spearhead region allows the former to ascend above through 370 Å, resulting in the uniform contraction of the tail sheath, as the upward thrust is generated from 6 equally spaced regions on the given plane, supported by LTFs and without any corkscrew type of upward motion. At this stage, the hub of the baseplate propped at a total distance of 900 Å from the host cell outer surface, the infection process reaches a metastable state. The credible reasoning in support of a uniform contraction and not a corkscrew type can be assigned to the absence of wrapped/twisted LTFs in the cryo TEM images depicting a P1 phage particle infecting its host, which would be necessary to rotate the baseplate hub for generating corkscrew type motion (SI 2) (1). The uniform contraction merely exposes a segment of the inner tube spearhead complex but increases the electronegativity in the inner sheath region resulting in further weakening of its grip on the tail tube. Subsequently, the entire phage transits axially downward in a corkscrew type of motion against the baseplate hub. The partially compressed sheath subunits suffer a non-uniform corkscrew contraction, originating from the top that drives the partially exposed tail tube attached to the spearhead complex downward to penetrate the host cell membrane and deliver the genome inside the host cell replication. The non-uniform sheath contraction would appear as a conical shape cylinder as seen and explained for T4 phage and R-type bacteriocins (7, 15). Due to the contraction, the sheath subunits rotate outward resulting in further increase of negative potential gradient to the interior of the sheath tube that would aid in the tail tube to move out of the system coupled with the downward corkscrew motion. The roles of the sheath and the tail tube along with the baseplate during the entire infection process are shown in Fig. 5B and Movie 1. Intermediates showing conical sheath forms, a part of the crock-screw motion, are scantily imaged in cryo/room temperature conditions as the phenomena has been suggested to be extremely rapid (Fig. S5B) (2). Owing to the lack of high-resolution structural information about the exact rearrangement of the baseplate proteins (especially gp16 and BplA^+-^ with PmgA) post-contraction, we have omitted that part in making Movie 1. It is interesting to note that each of the six protofilaments execute the non-uniform compression in unison thereby maintaining the six-fold symmetry of the sheath slabs during the entire infection process (Fig. S5C). The intermediate stages of the sheath compression reveal that the handshake motif of the sheath subunits is never compromised, nor the 6-fold symmetry of the sheath subunits show any ‘clash’ among each other during the dual contraction of the sheath subunits (Fig. 5B, Movies S6, S7). Maintaining the handshake mesh and therefore the six-fold symmetry of the gp22 horizontal slabs throughout sheath compression is the reason for the N and C terminus to gyrate differently than the rest of the gp22 molecule, as explained previously in this work.

**Fig. 5.**
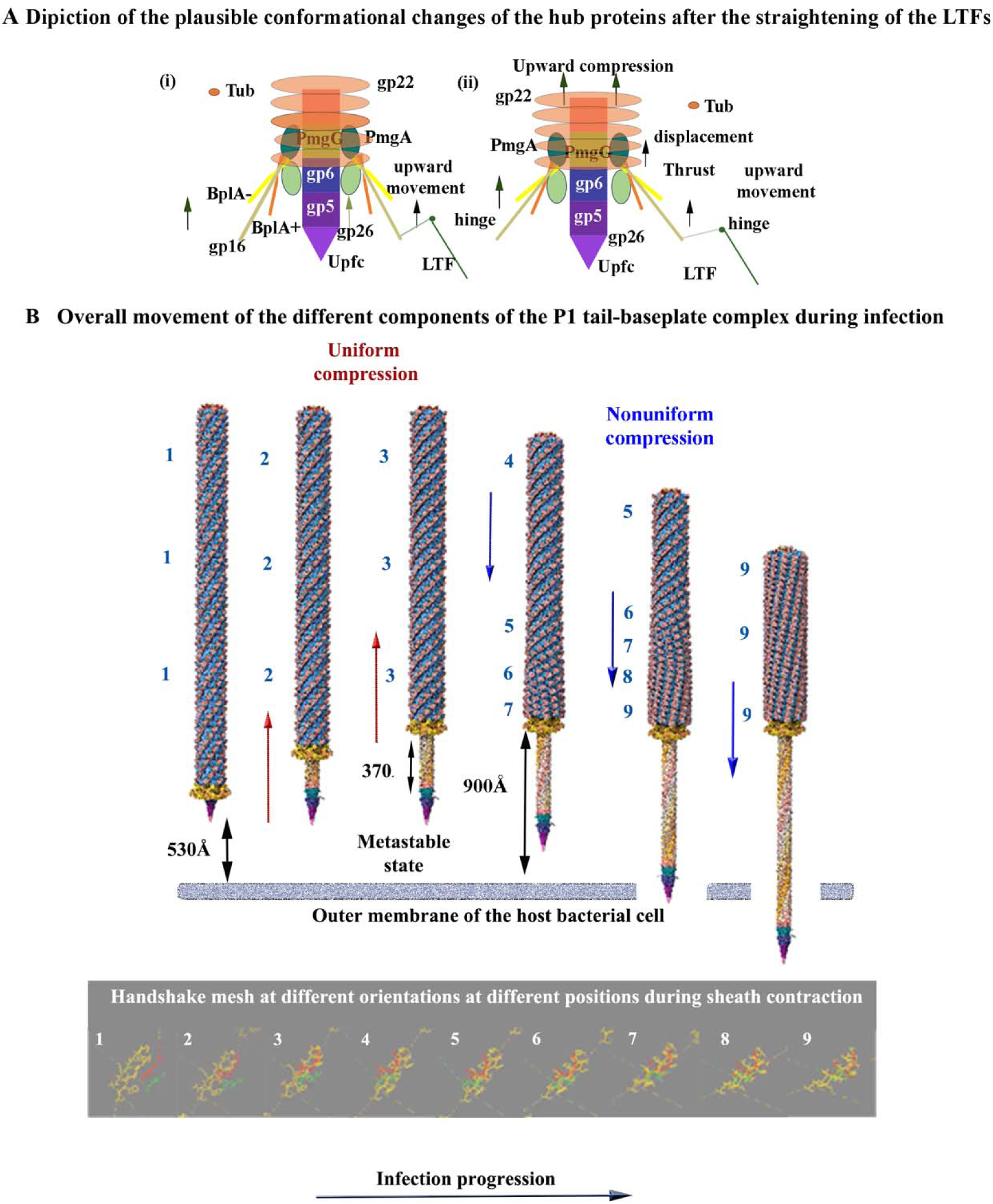
Action of the Pl virion tail-baseplate proteins during lytic infection process. (A) Schematic representation of the hub proteins before and after the straightening of the LTFs different proteins of the baseplate shown in (i) and (ii) respectively (summed up from Fig. 4D). The LTFs straighten up from their kink regions generating upward thrust resulting in the detachment of PmgA and gp26 from the spearhead region of the baseplate (ii). The transmitted upward thrust is utilized to compress the tail sheath in the upward direction. **(B)** Overall movement of the different components of the P1 TB complex during infection. The atomic structure of the TB complex of P1 phage is shown to be positioned at 530Å from the bottom line representing the host bacterial cell membrane. The arrows indicate the direction of the motion of the sheath and baseplate. Uniform upward movement of the hub region of the baseplate and sheath tube (due to the straightening of LTFs), exposing the spearhead portion of the baseplate and portion of the tail tube is followed by the non-uniform corkscrew downward contraction of the sheath followed by the downward motion of the tail tube puncturing the bacterial outer membrane and is summed up pictorially in Fig S5A. Each of the 6 states of the tail-baseplate complex is numbered (from 1-9). These positions are marked to show some of the changes in the protofilament orientations on the sheath tube due to the compressions. The orientations of the handshake mesh for those corresponding positions are shown below. The participating residues from three different subunits are Lime Green (15–18), Yellow (494-502 and 505-512) and Red (520-523), respectively.

## Discussion

Recent studies have reported that CISs can be successfully programmed to transport a medicinal cargo to desired targets in subcellular domains, employing their characteristic infection practice and have shown tremendous potential in treatment of dreadful diseases like cancer where conventional treatment processes have been found to fail (16). P1 phage, however, has a different mode of infection process in its lytic stage, as explained here. This study provides a guideline as where in P1, the desired payload can be devised so that it can be delivered to the target, employing the phage’s unique infection process which is primarily within the capsid and/or tail tube region. The use of minimalistic contact with its host would not require the P1 baseplate to diffuse through the biofilm matrix, making its attachment to the host redundant in the infectious process. Furthermore, it is recently suggested that the depolymerase found at the tip of the tail fibers of the phage disrupt the matrix of the biofilm leading to an unchallenging pathogenic host infection with great efficiency, supporting that the minimalistic contact of the P1 phage with its multiple pathogenic hosts is good enough to make a therapeutic effect is very much plausible that due to this minimalistic contact process, P1 can infect a variety pathogenic bacteria belonging to different strains, a characteristic unique to this CIS. Given that P1’s genetic mapping has been studied exhaustively, its ability to infect pathogenic bacterial strains of different species, coupled with results from this study, makes this virion ideal bio nano carrier to deliver therapeutic cargo to desired locations.

Another highlight of the dual compression is the length of the tail tube available to penetrate the host bacterial cell. The normal P1 sheath tube has a length of 2,100 Å, compresses to 1,150 Å, which attributes that 1,050 Å of effective penetrating rod comprised of tail tube + Tub + PmgG + gp6 (950 Å) and + baseplate needle (100 Å) descend 530 Å to reach the top of the outer membrane of the host cell, leaving 520 Å of the tail tube-spearhead complex, to penetrate through the host cell membrane to reach its cytoplasm to release the phage genome (Fig. S5A). This length is adequate to perforate through a normal *E* coli cell membrane (approximately 500 Å thick) to reach the cytoplasm. Cryo-ET work could visualize nearly 300 Å of tail tube penetrating the mini host cell, which stipulates that the entire spearhead region of length 230 Å (Tub + PmgG + gp6 + gp5 + UpfC) falls off after the penetration all the way to the cell cytoplasm. Similar observation has been made with the infection process of the T4 bacteriophage where the baseplate needle complex falls aside after punctuating the host cell, just prior to the transfer of the phage genome inside the host cell for replication.

The atomic structures of BplB, Tub, PmgG and gp6 possess a set of cross -strands forming their central inner sections that appear like a partial -barrel, an arrangement ideal for the transport of large protein complexes and this feature is well conserved for other CISs, irrespective of high or low sequence homogeneity (Fig. S6). Comparison of the atomic structures of the tail sheath and tube of P1 with other CISs show considerable similarities suggesting a common origin. All the 3 domains of gp22 show substantial structural similarity to that of phage E217, while domains 1 and 3 are in close proximity to the sheath subunit of R2 pyocin as domain 2 is missing in the later (4, 11) (Fig. S7A). Atomic structures of the tail tube subunits of P1 with E217 and R2 pyocin show significant similarities with the exception of the long 28 residues in the N-terminal region of the P1 (Fig. S7B). The tail tube subunit of P1 shows the best match with the T4 counterpart as both have long extended regions on the N-terminal domain. It is interesting to point out that the internal segment of the sheath tube has a -helix barrel like structure (discussed above) that is known to have high environment stability, a plausible reason that the tail sheath can withstand significant conformational changes yet maintaining 6-fold symmetry during the infection process (15, 17, 18). Our analysis reveals that each single protein of the TB complex, discussed here, has a unique and distinct role to play in the infection process and is not merely a structural protein and we are able to predict the possible roles of the proteins that have not shown in the density maps. In conclusion, the absence of STFs is the prime miscreant for this unique infection process, which is overcome by the retraction of the baseplate during infection and the dual contraction of the tail sheath leading to a successful infection process. The plausibility of dual contraction of the sheath subunits is supported in-parts by different myophages: Pseudomonas phage E217 supporting tail sheath upward contraction followed by T4 phage and R type bacteriocins for the corkscrew downward contraction during infection (11). We conclude by stating that CISs missing STFs would depend upon this unique exercise, like P1, for their biological functions.

## Supporting information

Movie S1. 3D view of the single gp22 subunit of the sheath of P1 phage showing the 4 domains with the start and end residues.

Movie S2. The gp22 molecule in its pre- and post- contracted states showing its gyration during the sheath tube contraction.

## Acknowledgements

The authors thank Dr. Tom Goddard of University of California San Francisco for his significant help in developing the movie showing the step infection process. The authors also thank Dr. Avery L. McIntosh and Dr. Lawrence Griffin of Texas A&M University for their support in this work. The authors also thank and acknowledge Mr. Gregory G. Servos of OvationData for his assistance with the project.

## Authors’ contribution

Conceptualization: AS. Methodology: AS. Specimen Preparation: AS and KK. Data collection: VB and AS. Computation image processing: AS, LT. Model building: TN, GT. Manuscript preparation: AS, TN, GT, DK, SC.

## Data Availability

Electron density maps for the sheath, tail tube, baseplate P1 bacteriophage have been submitted to the EMDB database with accession numbers as follows: EMD-27714 (pre-contracted sheath tube), EMD-27742 (post-contracted sheath tube), EMD-28019 (Tail Tube), EMD-43960 (baseplate of P1 phage).

PDB accession numbers: 9AY3(pre-contracted P1 sheath subunit), 9AY4 (Post-contracted P1 sheath subunit), 9AY2 (P1 Tail Tube subunit), 9AY5 (P1 baseplate).

## Supplementary Materials

Materials and Methods

Supplementary Text SI 1

Figs. S1 to S7

Movies S1 to S8

Supplementary Text SI 1

References

---

**Movie 1**: The atomic structure of the tail (tail tube, sheath, and baseplate) positioned at 530Å. As the LTFs push the baseplate upwards, the hub proteins undergo conformational changes, detach from the spearhead region of the baseplate, and ascend, resulting in the uniform contraction of the sheath till it is contracted by a length of 370Å. A ‘corkscrew’ type downward motion is then initiated which causes the sheath tube to compress non-uniformly, the tail tube attached to the spearhead region is driven downward and the infection takes place.

## Supplementary Materials for

### The PDF file includes

Supplementary figures Figs. S1 to S11

Materials and Methods

Supplementary Text SI 1

References 23 to 34

Supplementary Text SI 2

### Other Supplementary Materials for this manuscript include the following

Movies S1 to S8

Tables S1, S2

## Supplementary Figure Legends

**Fig, S1.**
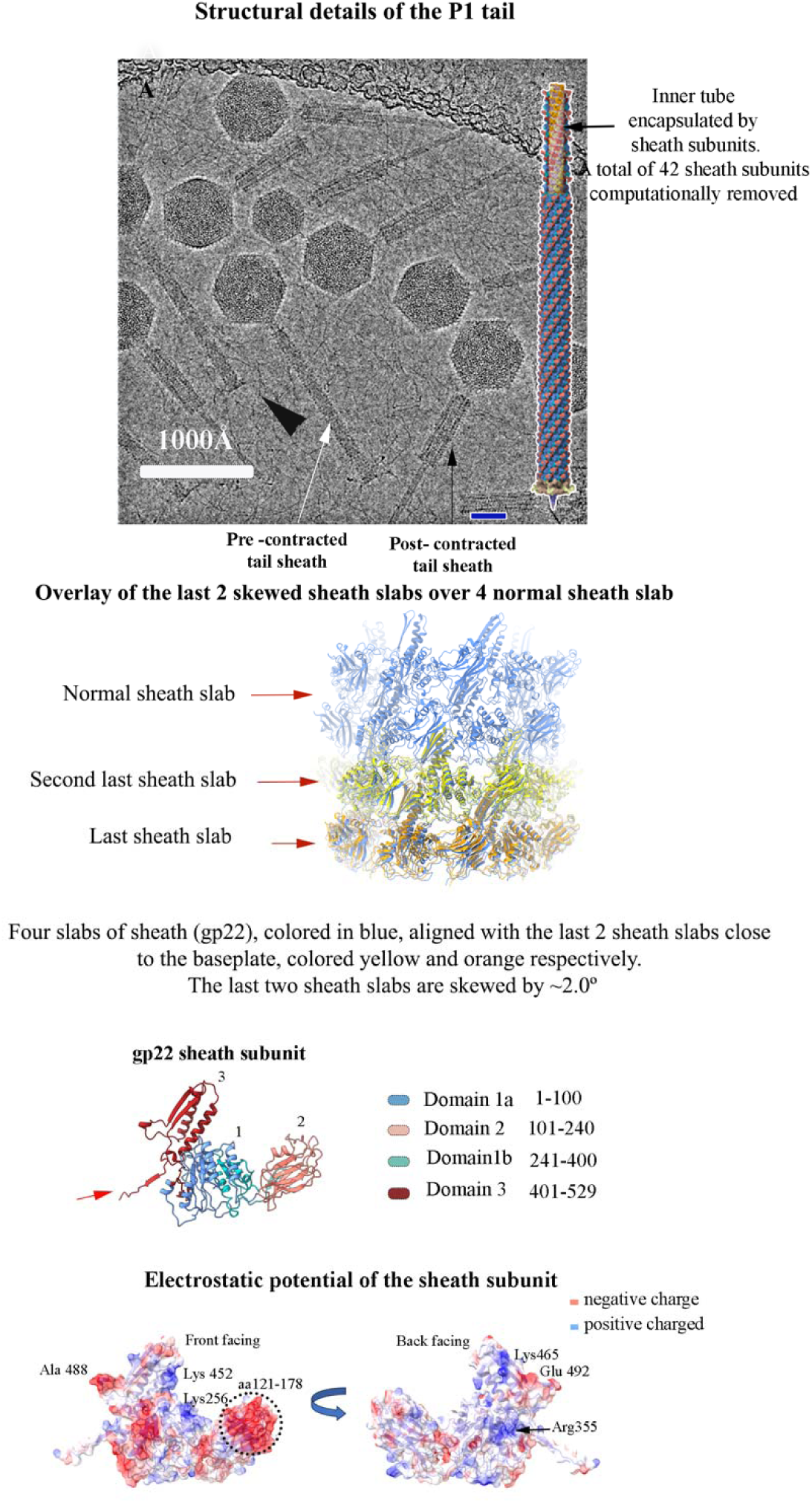
Cryo electron micrograph of P1 phages. White and black arrows indicate a P1 phage tail in its pre- and post-contracted states, respectively. The black arrow points to the LTFs. The black and grey arrows indicate the tail tube and the baseplate respectively. In the inset, 3D reconstructed tail of P1 phage with 48 sheath subunits computationally removed to show the positioning of the tail tube. Bar: 200Å. Overlap of the tail sheath subunits with the last 2 sheath slabs showing the skewing of the latter two by approximately 2.0° w.r.t the vertical axis. The 4 segments of gp22 are shown with color code and residues - Dark Turquoise 1-100, Salmon 101-240, Cornflower blue 241-400, Fire brick 401-529. Charged regions on the outer and inner surfaces of gp22. The positively charged residue (blue) residing in the interior of gp22 is responsible for holding the tail tube.

**Fig. S2.**
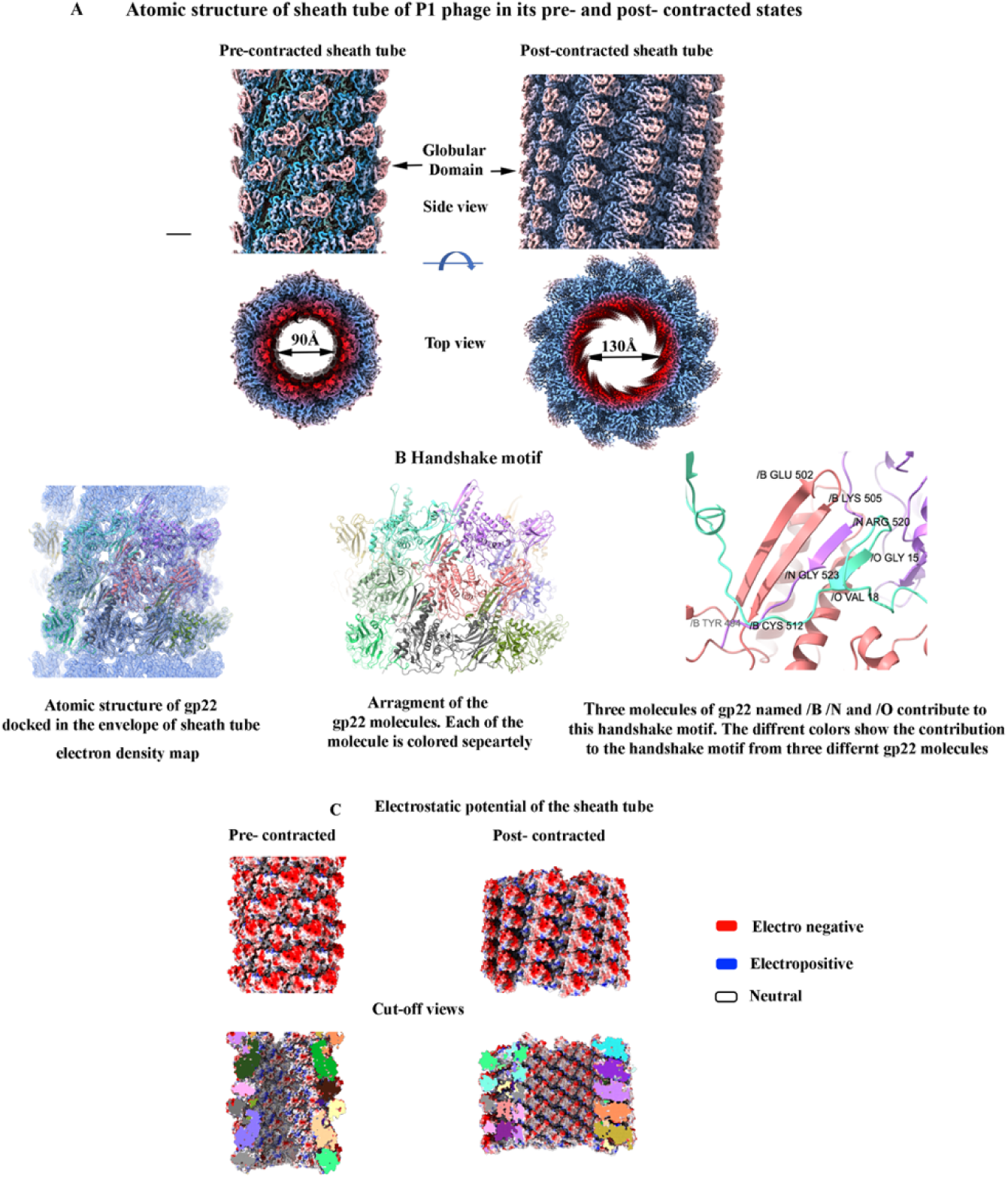
**(A)** Iso-surface representations of pre- and post-contracted electron density maps showing side and top views of sheath tube respectively. **(B)** Electron density map of sheath tube in its normal state with atomic structure of gp22 traced in the density envelope followed by the image of the atomic structure only (with the density map removed). Each of the gp22 molecules is colored separately. Zoomed in view of the handshake mesh, contributed 3 by gp22 molecules named /B, /N and /O respectively along with the residues. **(C)** Electrostatic surface of the inner and outer regions of the sheath tube in its pre- and post-contracted states. The negative charges (red) increase in the internal region after the contraction due to the rotation of the sheath subunits.

**Fig. S3.**
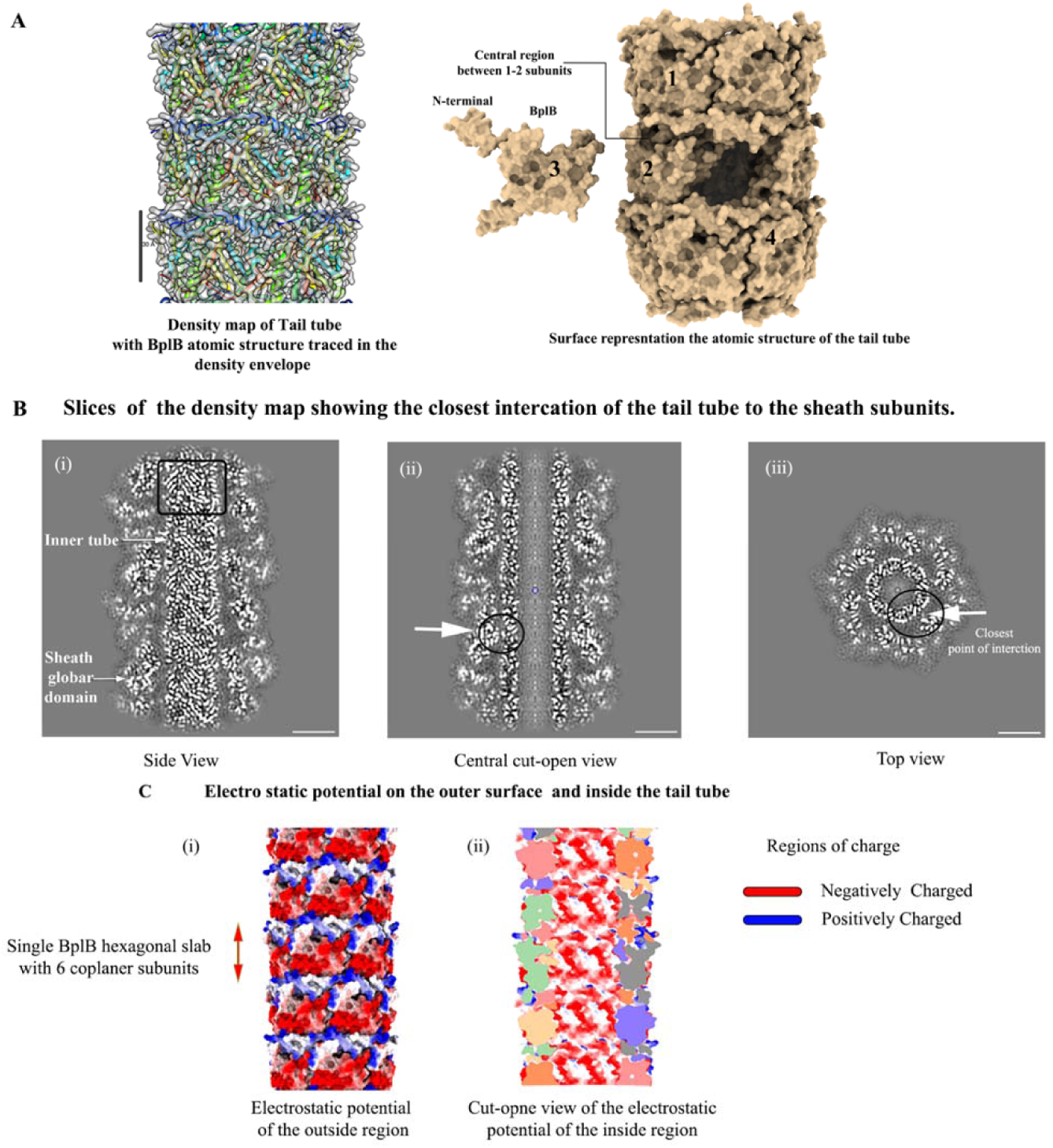
Electron Density map and atomic structure of Pl tail tube and its electro static potential. **(A)** Atomic structure traced in the 3D electron density envelope of the tail tube. Bar: 30Å. Surface representation of the atomic structure of the tail tube (BplB) with a total of 18 subunits of BplB group in 3 slabs, each with 6 subunits arranged in a C6 symmetry. The helical symmetry is the same as the tail sheath. A single subunit is pulled off to show the arrangement of the BplB subunits. **(B)** Grey scale slices at different positions visualized from the side (i, ii) and top (iii) of the tail electron density map showing the closest position of the tail sheath and tube, circled in black. **(C)** Electrostatic potential of the outer (i) and inner (ii) surface of the tail tube.

**Fig. S4.**
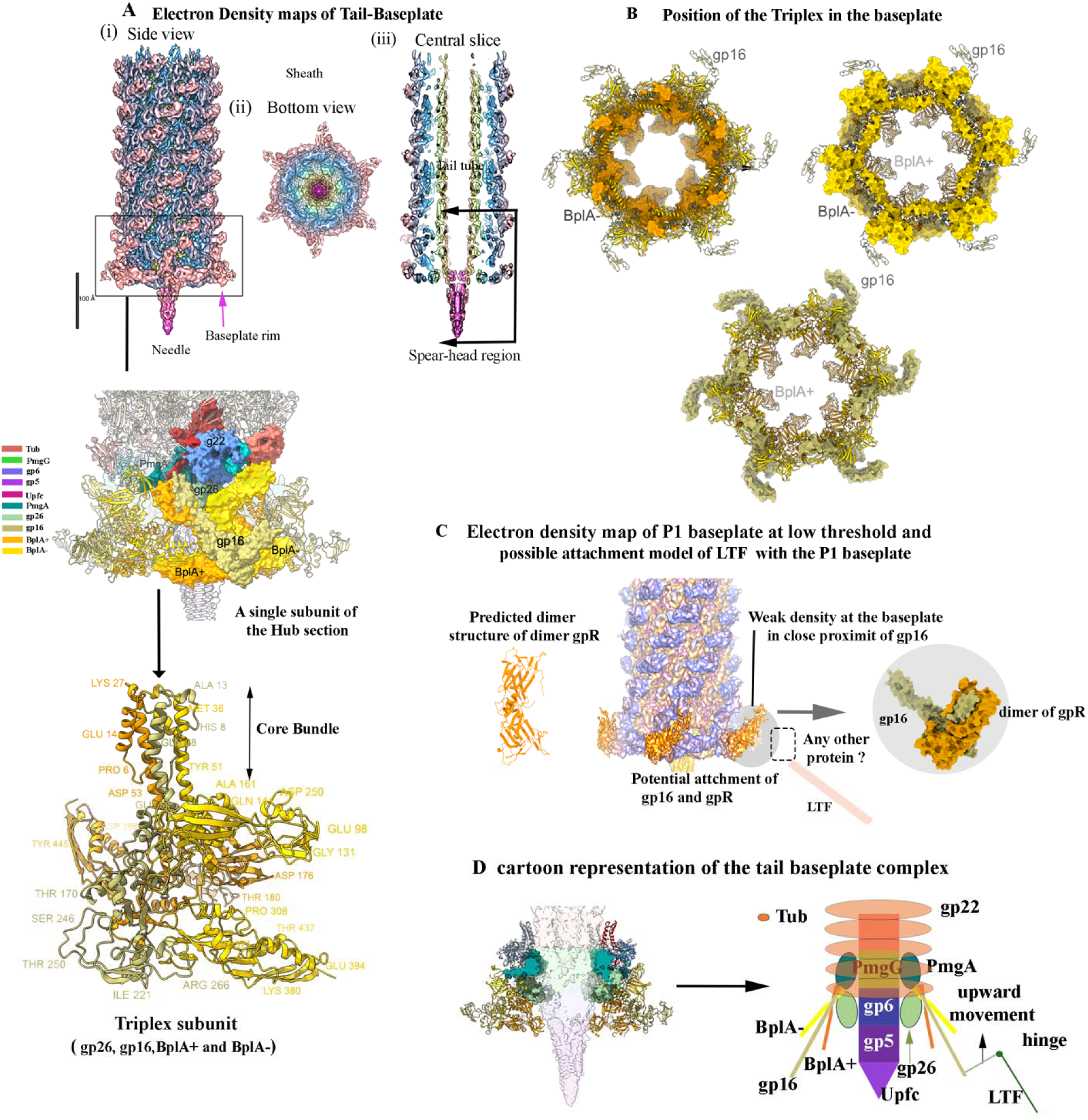
Organization of the different proteins forming the Pl baseplate. **(A)** Iso-surface representation of electron density map of the tail-baseplate region of P1 phage showing the side (i), and bottom view (ii). A central slice of the tail-baseplate complex showing the arrangement of the tail tube, the sheath and baseplate regions (iii). Bar: 100Å. Different proteins form the baseplate, primarily highlighting the baseplate hub along with a single gp22 molecule of the sheath from the last sheath slab with their identification colors. The triplex molecule is shown with some representative residues. Side view of a triplex molecule with some of the residues labelled following the color code. **(B)** Arrangement of the triplex molecules (BplA-, BplA+ and gp16). BplA+ is found to be holding the other two molecules BplA- and gp16. BplA-forms the outer rim of the baseplate while gp16 appears like a lever that brings in massive conformational changes with the initiation of the infection process (as explained in the text). **(C)** Section of the P1 tail-baseplate region shown at a very low threshold. The black elliptical circle shows the presence of very weak density, plausibly the point of contact with the proteins holding the LTFs. A predicted dimeric structure of gpR followed by a proposed model of gp16 and gpR dimer interaction. A potential attachment of LTF with the gpR and gp16 model is drawn. The box with dashed lines represents the region where unknown protein can reside holding the LTF to gpR. **(D)** A central section of the TB complex with a schematic representation of the same. This schematic is used in Fig. 5A to further explain the LTF-baseplate induced sheath contraction.

**Fig. S5.**
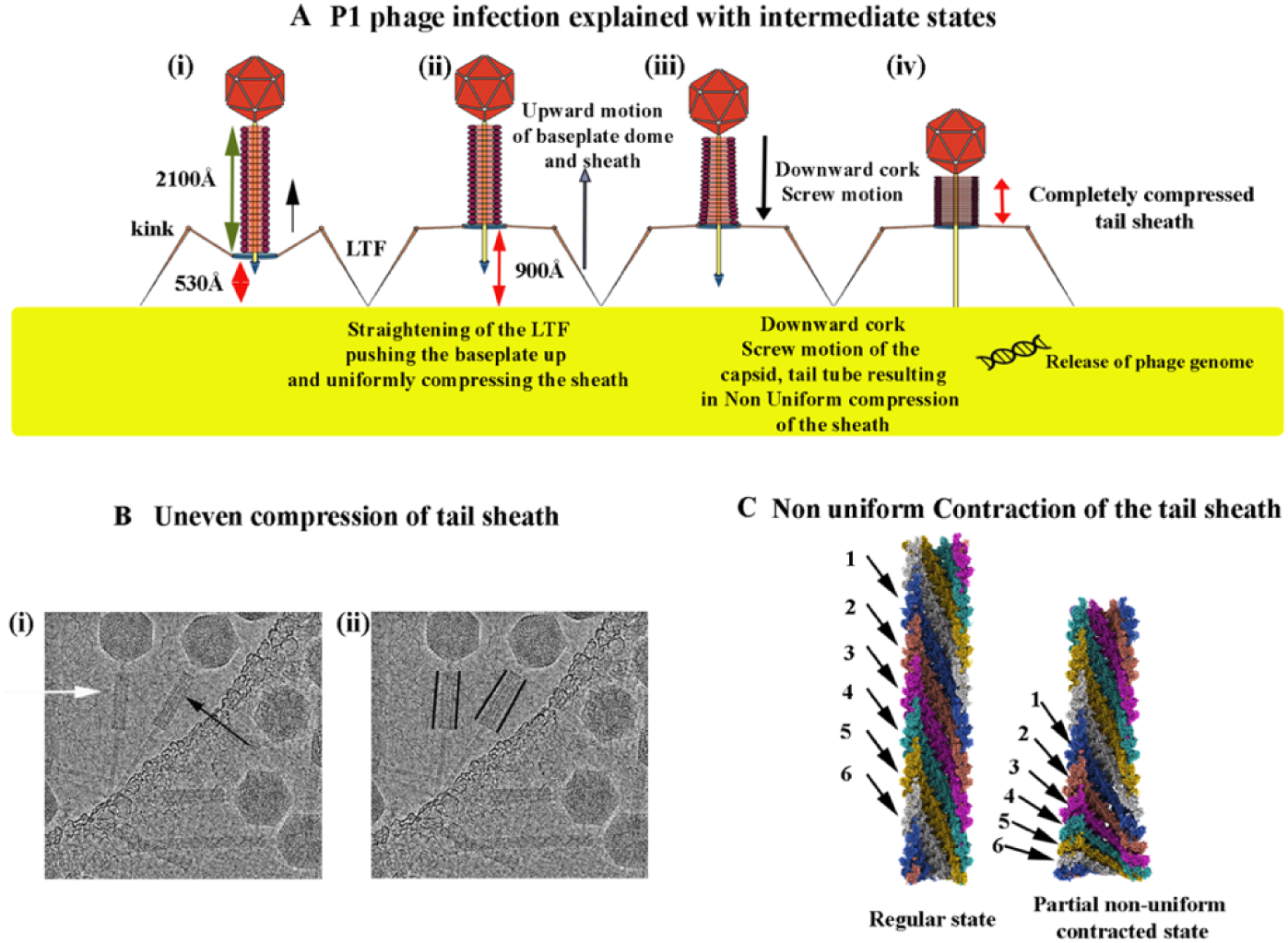
**(A)** Cartoon diagrams showing two step infection processes with the phage positioned on the host cell’s surface (i). Upward movement of the baseplate due to the straightening of the LTFs from their kink regions, acting as mechanical levers providing stability of the rest of the phage particle on the host cell surface and resulting in a uniform contraction of the sheath (ii) due to the straightening of the LTFs. Downward cork-screw motion resulting in a conical shape of the sheath and the downward movement of the tail tube toward the host cell (iii). Complete contraction of the tail sheath with the tail tube puncturing the host cell membrane (iv). **(B)** Cryo image of two P1 phage with contracted tail sheaths. The phage particle pointed out by a white arrow shows a uniform contracted sheath while the second, pointed by a black arrow has a slightly conical shaped contracted sheath suggesting a cork-screw type motion (i). Two black lines bordering the outlines of the contracted sheaths of the two phage particles to enhance the comparison between their shapes (ii). **(C)** Cut-open view of the Uniform and non -unform compressed sections of the tail sheath. A total of 6 protofilaments colored differently are pointed by black arrows and numbered accordingly. The cut-open views of the two states enables us to understand how each of the sheath slabs behave differently from each other under a corkscrew motion, with non-uniform diameters expanding outward, yet maintaining the interconnectivity among each other.

**Fig. S6.**
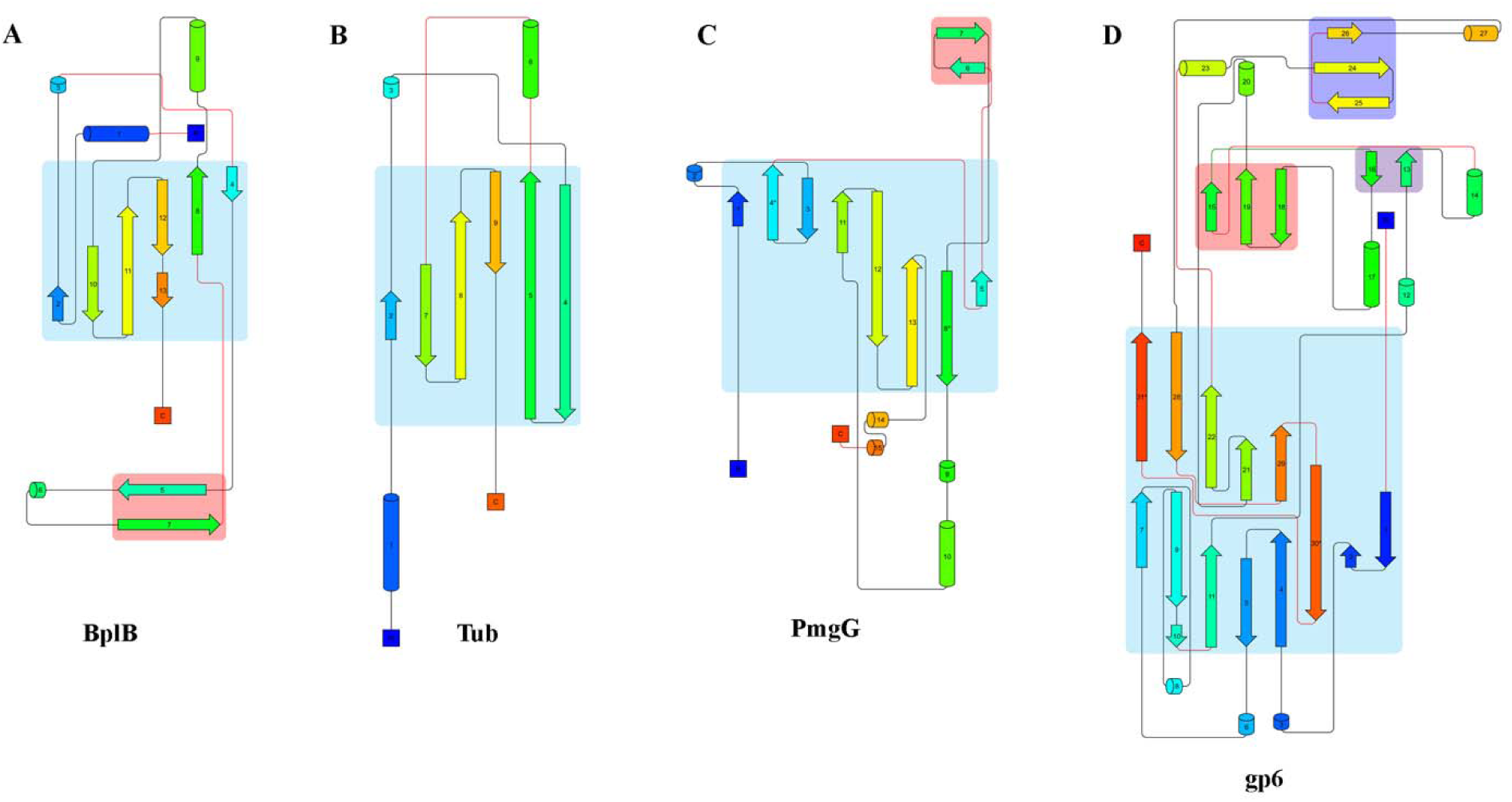
Topological diagram of different proteins forming tell central section of the tail. The blue rectangles show the set of strands forming the central section of the subunits. The cylinder represents helix and arrow represents a stand.

**Fig. S7.**
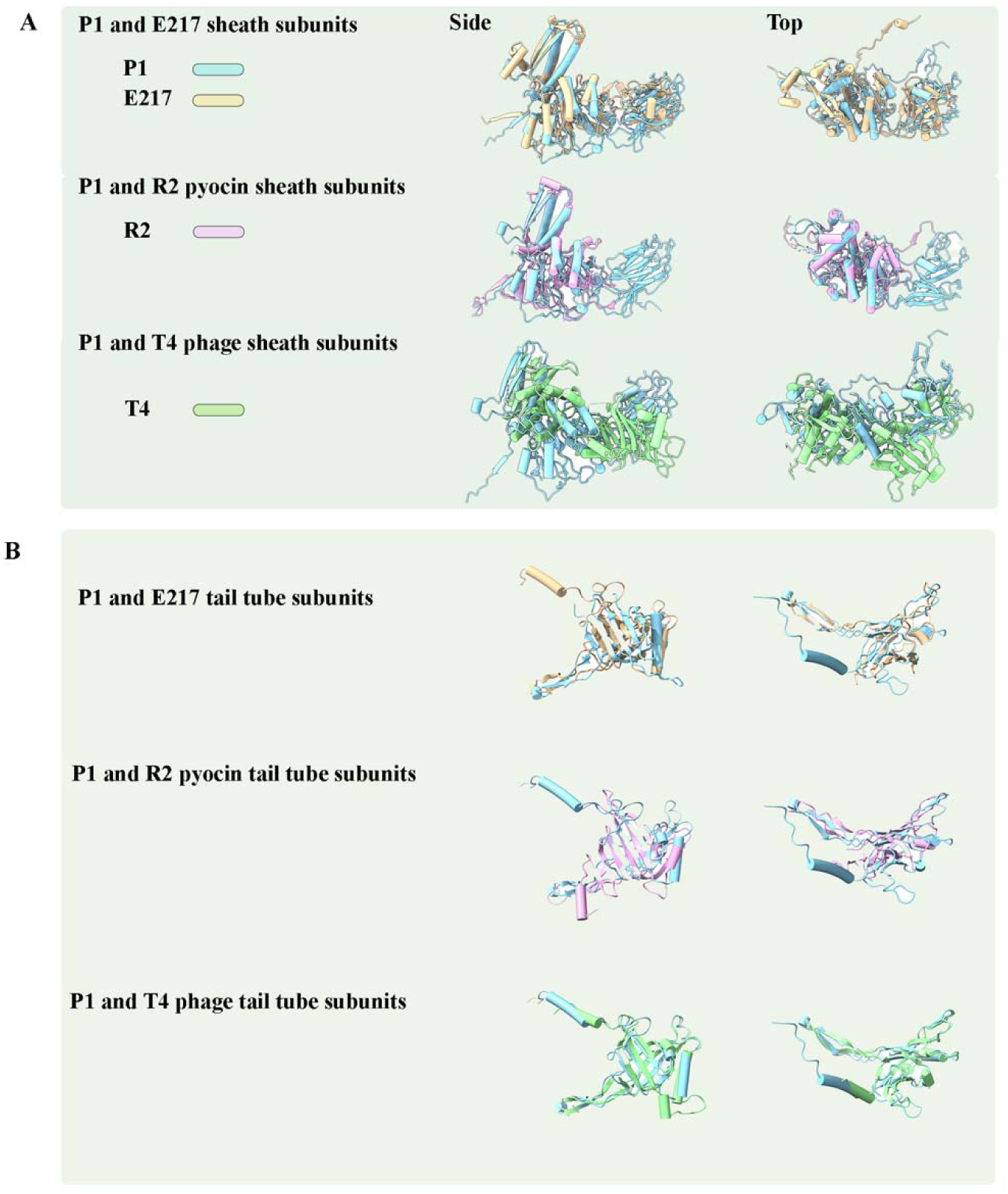
The sheath **(A)** and tail tube subunits **(B)** of P1 are overlaid with those of E217, R2 pyocin and T4 bacteriophage. The side and the top views reveal that the tail tube subunits of all the four CISs (E217, R2 pyocin and T4) are very much like each other while their sheath subunits show some less resemblance.

## Materials and Methods

### Virus Propagation and Purification

Bacteriophage P1 (ATCC 25404-B1) was propagated using a host bacterium Escherichia coli following a confluent lysis method (23). The host was grown in Luria-Bertani (LB) broth (10 g/L tryptone, 5 g/L yeast extract, 10 g/L NaCl, and 10 mM MgSO4) until its optical density at 600 nm wavelength (OD600) reached 0.35 (HACH DR6000), which was then inoculated with P1 phages at multiplicity of infection of 1 in presence of 5 mM CaCl2. A 100 µL of the mixture was added to 10 mL soft LB agar (LB broth amended with 5 g/L agar and 5 mM CaCl_2_) and poured on a LB agar plate (LB broth amended with 15 g/L agar and 5 mM CaCl2). The plate was incubated at 37 °C until a clear phage lawn appeared because of confluent lysis. The lawn was collected and centrifuged (10,000 g for 10 min at 4 °C) after which supernatant was recovered. For further purification, supernatant was filtered with 0.45 µm polyethersulfone (PES), pelletized (15,000 g for 24 hours at 4 °C), and resuspended in 3 mL SM buffer (50 mM Tris-HCl, 100 mM NaCl, 8 mM MgSO4, pH 7.3) overnight at 4 °C. Afterwards, the resultant was passed through CsCl layers (1.3-1.6 g/cm3 prepared in SM buffer, 104,000 g for 24 hours at 4 °C). Phage band was recovered, pelletized (15,000 g for 24 hours at 4 °C), and resuspended in 1 mL SM buffer overnight at 4 °C. The titer of the final stock was found to be approximately 10^11^ PFU/mL based on the double agar layer method.

### Sample preparation for cryo-TEM and data collection

Copper 300-mesh grids supported with a formvar layer and coated with an ultrathin layer of carbon were prepared by glow-discharging for 1 min at 10 mA under vacuum (PELCO 118 easyGlow). About 5μlts of P1 virus stock (10^12^ particles/ml) suspension were directly mounted on the grids followed by negative staining with 2% uranyl acetate solution (pH not adjusted). Grids were screened using ThermoFisher TF20 operating at an accelerating voltage of 200 keV equipped with Gatan K2 direct detector and Gatan Tridiem GIF-CCD camera for quality and concentration of the virion particles.

For cryo-TEM, approximately 3.0 µL of purified virus stock was loaded on holey carbon film grids with 2.0 µm holes (C-flatTM 124, Electron Microscopy Sciences), blotted in an environmentally controlled chamber at 80% humidity, and vitrified by sudden plunging the grids into liquid ethane at -180 °C (EM GP2 Automatic Plunge Freezer, Leica). Prepared cryo-samples were imaged with ThermoFisher TF20 operating at an accelerating voltage of 200 keV equipped with Gatan K2 direct detector, Gatan Tridiem GIF-CCD camera and ThermoFisher Glacios cryo-TEM equipped with an autoloader and Falcon 4 direct electron detector (DED). The grids carrying the vitrified sample were imaged on Glacios cryo-transmission electron microscope (ThermoFisher Scientific) equipped with a field emission gun operated at 200 kV. The microscope was aligned at parallel illumination. Grids were screened for ice quality and particle distribution. For the most promising grids, a total of 4200 movies were recorded in EER format using a Falcon 4 DED at a nominal magnification of 73,000X that corresponded to a calibrated physical pixel size 1.378 Å/pixel (24). The movies were acquired within a defocus range of -0.5 μm to -1.5 μm in an automated fashion using EPU software (ThermoFisher Scientific). Each movie was recorded in counting mode using a total exposure dose of 30 e/Å^2^ at a low dose rate of 3.2 e/Å^2^/sec to avoid coincidence loss. Additional details of data acquisition and processing are summarized below.

**Table S1.** Glacios cryo-TEM set up and data collection parameters.

|  |  |
| --- | --- |
| Nominal magnification | 73,000x |
| Apertures ($\mu\text{m}$) | C2 aperture = 50 $\mu\text{m}$; Objective aperture: none |
| Spot size | 5 |
| Pixel size ($\text{\AA}/\text{pixel}$) | 1.3889 |
| Total dose (e-/Å<sup>2</sup>) | 30 |
| Total number of EER frames per movie | 2,267 |
| Exposure time (sec) | 9.4 |
| Defocus | -0.5 to -1.5 μm |

### Computational Image processing

The frames of the movies of the TF20 cryo data recorded were aligned using MotionCor2 (25). Movies with frames that showed poor alignment were left out. About 6400 and 5800 overlapping boxes for pre- and post-contracted tail states were generated and their power spectrums were computed with values of the reflections shown (Figure S8). The helical parameters of the pre- and post-contracted states of the tail sheath were estimated: the axial rise (Δx) is estimated at ∼39.0 Å and ∼17.0 Å, respectively. The rise angles (Δ∅) for pre- and post-contracted states were estimated to be ∼19.0° and ∼30.2° respectively. Computational image reconstruction of the pre- and post-contracted states were carried out using bhelix of BSOFT package with a featureless solid cylinder a starting model and maps were obtained to a resolution of ∼ 13.0 Å at an FSC cutoff of 0.143 (26). These maps were used as starting models for high resolution maps.

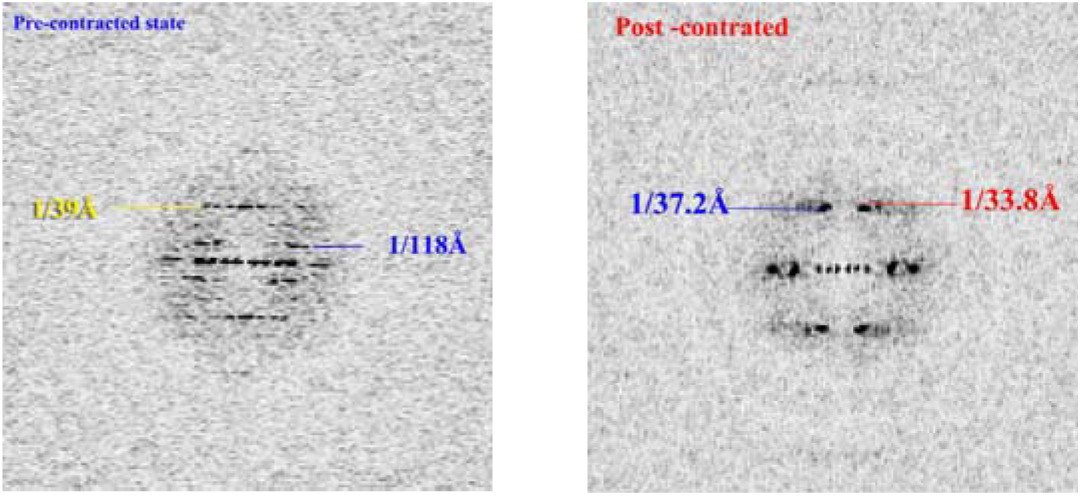

The movies collected from 200 keV Glacios cryo-TEM were fed in the cryoSPARC software package and frames were aligned using Patch Motion Correction (27). The aligned frame providing one micrograph for each movie were utilized for CTF estimation using Patch CTF. Overlapping boxes of the pre- and post-contracted state of the tail of P1 were generated followed by 2D class averaging and the best class averages were chosen to be used as a template for further auto boxing more segments of the tail section. Another 3 rounds of 2D class averaging were carried out to select the best particles for reconstruction. The 3D maps computed earlier using BSOFT were employed as initial reference maps, and the final reconstructions were carried out using the boxed particles selected after a few rounds of iterations were carried out. For the tail baseplate reconstruction, a similar approach was taken, and the reconstructions were carried out in CryoSPARC (27). The resolution of the different density maps at Fourier Shell Correlation (fsc) cut-off of 0.143 is shown below. The density maps were visualized using UCSF Chimera and ChimeraX (28). The movies are made in ChimeraX.

**Table S2.**
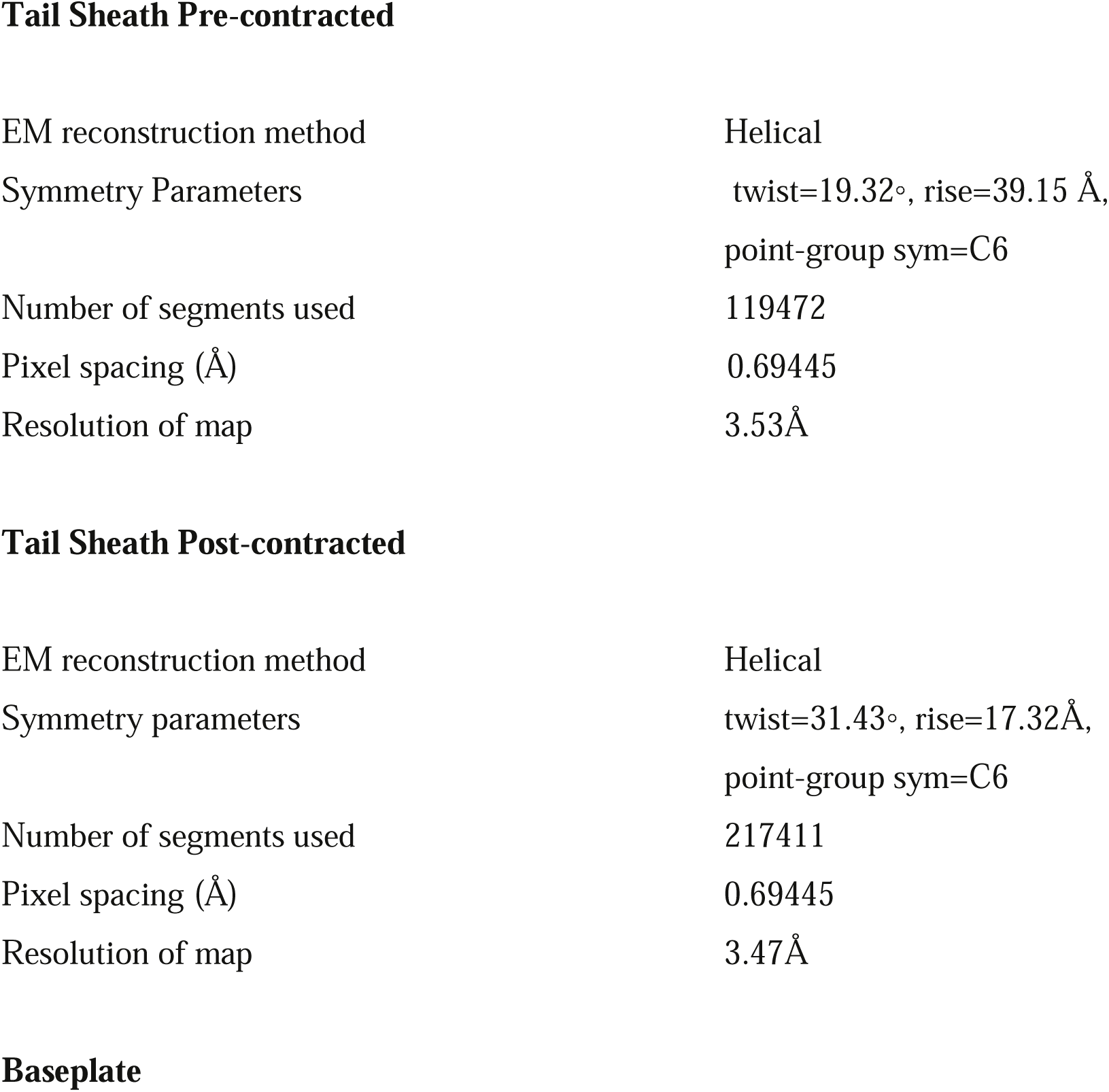
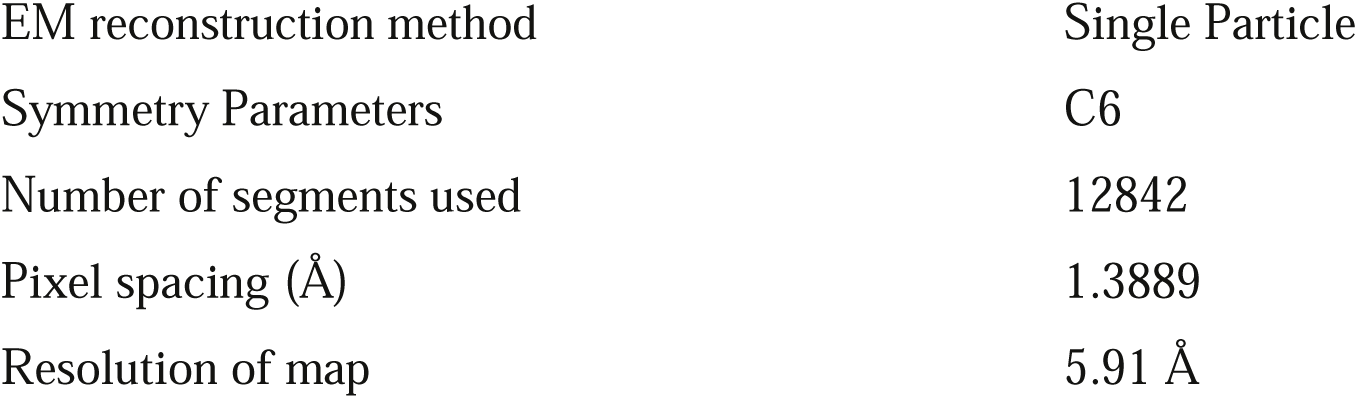
Statistics for reconstruction.

**Tail Sheath Pre-contracted**
|  |  |
| --- | --- |
| EM reconstruction method | Helical |
| Symmetry Parameters | twist=19.32°, rise=39.15 Å,<br>point-group sym=C6 |
| Number of segments used | 119472 |
| Pixel spacing (Å) | 0.69445 |
| Resolution of map | 3.53 Å |
| **Tail Sheath Post-contracted** |  |
| EM reconstruction method | Helical |
| Symmetry parameters | twist=31.43°, rise=17.32 Å,<br>point-group sym=C6 |
| Number of segments used | 217411 |
| Pixel spacing (Å) | 0.69445 |
| Resolution of map | 3.47 Å |
| **Baseplate** |  |
| EM reconstruction method | Single Particle |
| Symmetry Parameters | C6 |
| Number of segments used | 12842 |
| Pixel spacing (Å) | 1.3889 |
| Resolution of map | 5.91 Å |

**Figure S9.**
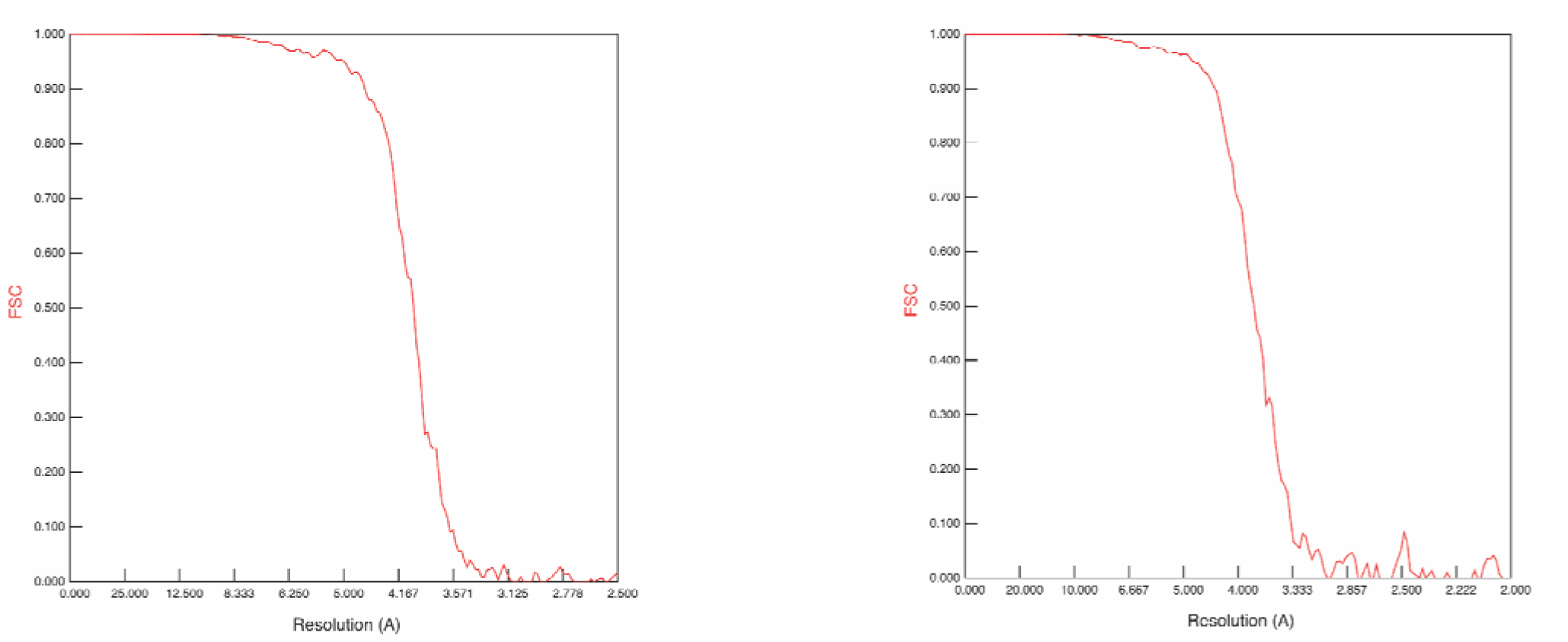
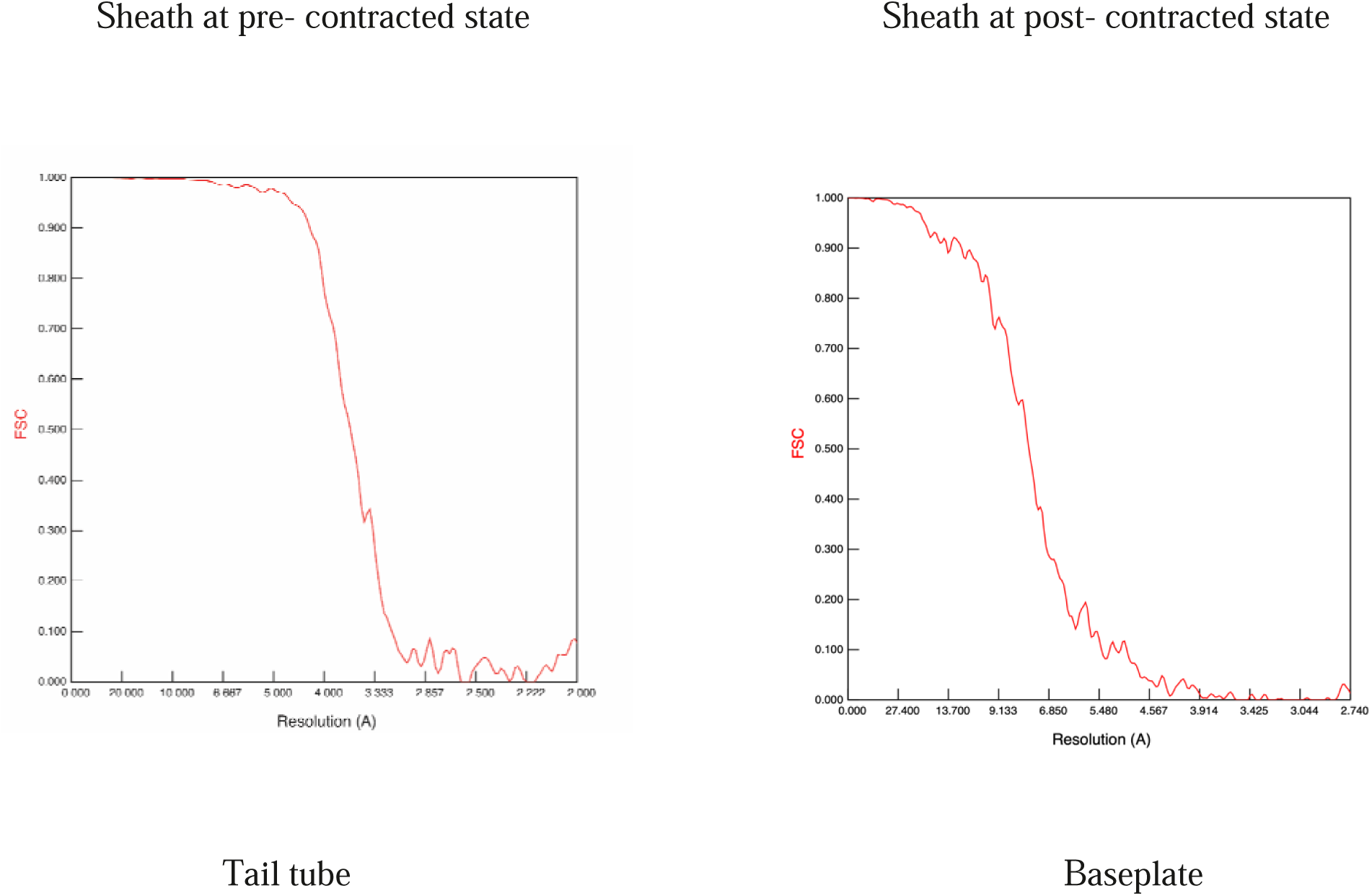
Resolution curves of the density maps of the different section of P1phage tail.

Resolution determination method - Resolutions of the maps were estimated in cryoSPARC with gold-standard Fourier shell correlation (fsc) at a cutoff value of 0.143 with high resolution noise substitution. The tail tube region is segmented out of the density map of the pre-contracted tail. The resolution of the tail tube is estimated from the segmented regions from the pre-contracted tail sheath half maps. All the half maps of different segments of the P1 tail were masked to determine the resolution of their corresponding final map.

### Computing helical parameters of the tail sheath at the metastable state for building the Movie 1

During the infection process, the P1 baseplate is found to have receded through ∼370 Å (1). Following the dual compression infection process explained here in this work, the tail sheath would subsequently compress through 370 Å, the partial contracted state, (can also be called a the metastable state) of the helical sheath would have a length of ∼1730Å. There are ∼53 slabs of 6-fold symmetry sheath subunits for the entire sheath tube. So, the axial rise of the helical sheath subunit at this partial state would be ∼32.6 Å. As the initial contraction of the tail sheath is considered a uniform throughout the tail sheath tube, reaching the partial contracted state, the turn angle for that state can be computed from the plot below (using the helical parameters of non-contracted and fully contracted sheath (Table S2). The value of the twist angle with an axial rise of 32.6 Å is ∼22.6°.

**Fig. S10.**
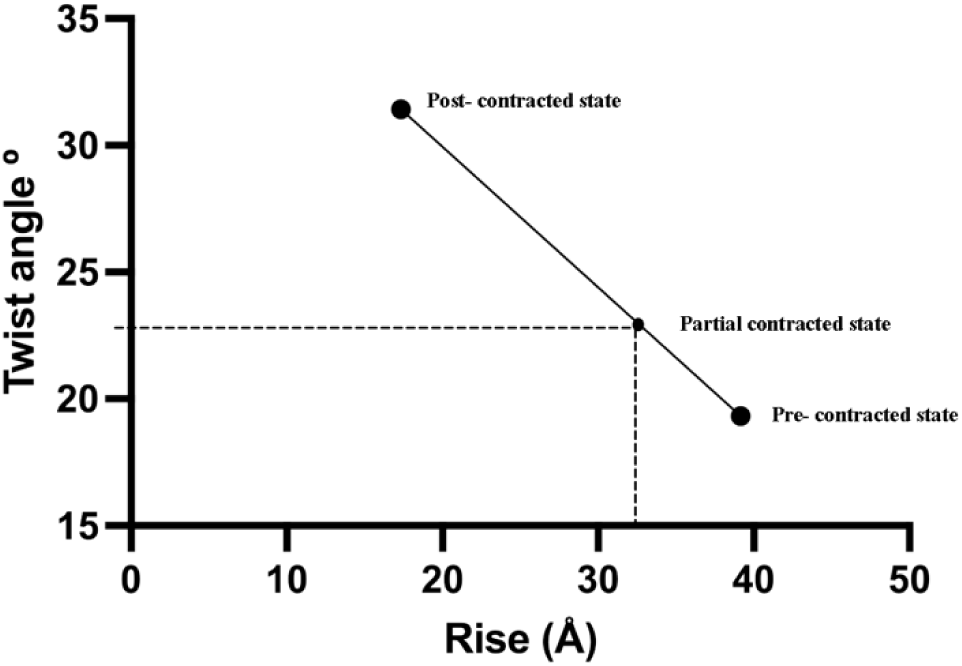
Plot of the helical parameters of P1 sheath in pre-and post-contracted states.

### Model building and refinement

We used AlphaFold2 to generate initial predicted atomic models for the P1 phage proteins listed in Table 1 (29). These predicted models were then manually or automatically fitted into the EM map using UCSF ChimeraX (28). For the sheath and innertube, DeepMainmast was also used to build atomic models (30). DeepMainmast employs deep learning to capture the local map features of amino acids and atoms to assist main chain tracing and can consider AlphaFold2 models (30). For the baseplate, to determine which monomers should be predicted together, we referred to the T4 phage structure (PDB-ID: 5IV5) (2). Fitted models were manually adjusted and rebuilt using Coot (31). For model refinement, we performed Rosetta relax protocol and real-space refinement in Phenix package (32, 33). We performed multiple iterations of model fitting and refinement to improve the quality of the models. To verify the quality of the models, we used Comprehensive validation tool in Phenix, and a map-model quality assessment method called DAQ score (34). For the chains modeled on maps having better than 5 Å resolution, the assessment results by DAQ score are shown in Fig. SI.4. The following section describes the details of the modeling for each component in the P1 phage.

### Sheath

For the sheath structure model, we used two EM maps corresponding to two structural states, pre-contracted and post-contracted, respectively. The initial predicted atomic models of the sheath proteins were built using AlphaFold2 with the gp22 sequence (Table 1). The gp22 predicted models of the monomer and 2-6 homo-oligomer models were constructed with AlphaFold2, and for each EM map, the structure model that fits the EM map the best was selected for further model-building steps. During the iterative refinement phase of the sheath model, a single gp22 protein predicted model was refined along with surrounding chains without non-crystallographic symmetry (NCS) constraints and restraints. In the pre-contracted model, a predicted model was refined with eight surrounding chains, while in the post-contracted model, nine surrounding chains were used for refinement. The refined chain was then copied in symmetric positions to make the sheath tube structure. To enhance the fitness of the gp22 ring models to the EM map while maintaining the symmetry of the structural model as much as possible, refinement was performed in the order of with NCS constraints, with NCS restraints, and without NCS. To avoid forcing the same copy between ring models and reducing the effect of erroneous refinement from both ends of the tube, the NCS groups for each hexameric ring were manually defined. Then we went back to the step to refine a monomer chain model and repeated this refinement round several times. Finally, the auxiliary rings, which acted as interacting pairs during refinement, were then removed from the top and bottom of the sheath tube. Pre-contracted chains were refined as five rings and finalized with the middle three rings, considering a chain is adjacent to the other chains on the upper and lower rings. Similarly, post-contracted chains were refined as nine rings and finalized with the middle five rings, considering that a chain is adjacent to the other chains on the upper and lower two rings. In the sheath protein structure of both pre-contracted and post-contracted states, gp22 proteins were connected by a “handshake” domain that consists of two antiparallel b strands (strand1 and strand2, aa 494-499, 506-512) of one gp22, one parallel b strand (strand3, aa 521-527) of a neighboring gp22 and one parallel b strand (strand4, aa16-18) of another neighboring gp22 on the upper layer (see Fig. S1C, S2B). Since AlphaFold2 could not build the handshake domain, especially two parallel - strands (strands 3 and 4), we used Coot to optimize the orientation of the side chains and adjust the backbone structure. The constructed models were evaluated with DAQ scores (Fig. SI.4(a,b)).

### Innertube

We used AlphaFold2 to build predicted atomic models of BplB and Tub for the innertube structure. After comparing both models, we selected the BplB model for the innertube due to its much better fitness of loop regions (aa103-108 and aa113-130) to the map. During the refinement of the inner tube model, we followed the same protocol as for the sheath model’s refinement step. The model was refined as five rings and finalized with the middle three rings, considering a chain is adjacent to the other chains from the upper and lower rings. Fig. SI.4.(c) shows the assessment results by DAQ scores. The BplB model has positive DAQ(AA) scores for almost all regions. The region (aa115-117) has low DAQ (Cα) scores around –0.5, which is due to the lack of volume for the middle G121 of S120, G121, V122, possibly causing an odd input shape of the volume to DAQ. Also, DAQ scores for a predicted Tub monomer model were calculated. As shown in Figure S11 (d), DAQ scores are overall low indicating that the Tub model does not fit the innertube map. The Tub was predicted as a hexamer and superimposed as a hexamer on the volume manually similar orientation to refined BplB.

### Baseplate

It was found that the last bottom part of the volume of innertube in the baseplate did not support the gp22 model well. Therefore, the Tub hexameric model was placed one ring below the gp22 innertube rings. 23 residues from the N-terminal of the Tub model were removed due to the absence of the corresponding volume in the map. Additionally, the PmgG hexamer model was placed at the last hexameric symmetry part of the inner baseplate based on the sequence similarity of P1 PmgG with T4 gp48 (5IV5 chain U).

For the wedge composing proteins, we first selected BplA based on the sequence similarity of P1 BplA with T4 gp6 (5IV5 chain A). Next, we built a trimer structure model that consisted of the BplA dimer (BplA+, BplA-) and gp16 monomer, whose function was unknown previously (22). The reason for selecting gp16 for the remaining volume in the wedge composing protein is because gp16 is next to BplA in the same operon in the P1 genome, and there are weak sequence similarities between the N-terminal (res. 38-64) of gp16 and N-terminal (res. 51-78) of BplA. Additionally, results from extensive predictions using Alphafold2 in terms of combinations of proteins remaining the possibility of working as the wedge and potential copy numbers suggested that gp16 was the best candidate. Based on the sequence similarity between P1 gp26 and T4 gp53 (5IV5 chain V), as well as between PmgA and T4 gp25 (5IV5 chain T), the remaining volume was filled with models of gp26 and PmgA. The Sheath gp22 ring model in the baseplate was refined considering NCS between chains in the same ring, like the refinement of the sheath. This refinement procedure enables us to observe that the last two rings are skewed, as shown in Fig. S1E. The other components were also refined as well.

### Spike

Gp5 trimer, gp6 trimer, and UpfC monomer model were predicted as one multimer using AlphaFold2. The predicted model was superimposed without refinement onto the map obtained from three-fold symmetric reconstruction.

### gpR & gp16

We observed that there was a blurred volume at the edge of the wedge. Based on the sequence similarity between P1 gpR and Pecentumvirus A511 gp105 (6HHK), which is a known T4 gp8 (5IV5 chain D) ortholog (12). Though predicted gpR dimer and gp16 showed high structural similarity with the counterparts of T4 (5IV5 chain D, E, and C (T4 gp8, gp8, and gp7)) and overall predicted model shape was relatively matched with the remaining volume, the volume was not clear enough to support the predicted model and proceed to refinement. Thus, we just made and placed a chimera model by combining a partial model of refined gp16 from the wedge with a partial model of predicted gp16 that attaches to the gpR dimer. In this process, the orientation between two gp16 models was adjusted to place the gpR dimer in the remaining volume.

## DAQ scores for a chain from each component

**Figure S11.** DAQ scores for a chain from each component modeled on a map having better than 5Å resolution. All calculations were for chain A for each component. The DAQ(AA) and DAQ(Cα) scores are shown along the sequence in plots. In general, a residue in a model has a positive score for DAQ(AA)/DAQ(Cα) if the amino acid/Cα position assignment is likely to be correct. The horizontal red lines are on three cutoff values at 0.0, -0.5, and -1.0. (d) DAQ score for attempting placing Tub on innertube.

## Supplementary information (SI) 2

### Reason for dual compression, explained

There are two distinct motions involved during the infection process of P1 phage. One is the upward movement of the ‘hub’ of the baseplate, in the direction away from the outer membrane of the host cell and could only happen if the LTFs straightened upward, like levers, from their kink region, propelling the hub up, as suggested by the cryo-ET work (1). This is a consequence of the absence of the STFs (established in this current work), that are necessary for the baseplate to clinch to the host’s outer surface in proximity, as seen in T4 bacteriophage (2). The second is the downward corkscrew motion of the phage capsid, the neck region, the tail tube and the spearhead region penetrating the host cell membrane for infection (2, 7). It is evident from the structural organization of the gp22 protein subunits forming the sheath tube that two opposing motions cannot act at the same time to maintain the structural integrity (C6 in-plane symmetry) of the sheath tube and therefore can be established that only one motion takes place at a given time.

After the attachment of the LTFs to the outer surface of the host cell, a corkscrew downward motion of the P1 phage would lead to a successful penetration of the host cell membrane like a typical myophage infection, without the need for an upward motion of the baseplate hub. Moreover, the upward movement would push the phage particle away from the host cell surface, something unfavorable for the phage infection process. This is a violation of the results seen in the P1 infection process using Cryo-ET (1). Thus, we conclude with confidence that the downward corkscrew motion is not the initial but the upward motion of the baseplate hub to be executed first by the P1 phage during infection.

Cryo-ET reports that the baseplate hub travels 370Å up and away from the host cell outer surface indicating that there is a partial compression of the tail sheath which has a length of 2100Å. This partial compression of the sheath tube cannot be an upward corkscrew motion for the following reasons. If the baseplate hub performs an upward corkscrew motion, the sheath would form a partially contracted semi-conical shaped sheath tube and would encounter a downward corkscrew motion next, a necessary act for the infection process (2, 7). That would disrupt the structural symmetry of the sheath tube and would result in a failed infection process. Furthermore, for the baseplate hub to perform an upward corkscrew motion the LTFs must twist themselves. However, no such LTFs in ‘twisted’ states were imaged in the cryo-ET (1).

Following the above discussion, we sum up the sequence of the two motions. As LTFs attach to the outer surface of the host cell and start straightening, the baseplate hub detaches from the spearhead region and ascends resulting in a partial uniform compression of the sheath positioning itself at 900Å away from the outer surface of the host cell. The second motion commences that causes the partially compressed sheath to suffer a non-uniform downward compression and would lead the entire phage except the baseplate hub to descend in a corkscrew manner for the infection.

## Movies

**Movie S1**. 3D view of the single gp22 subunit of the sheath of P1 phage showing the 4 domains with the start and end residues.

**Movie S2.** The gp22 molecule in its pre- and post-contracted states showing its gyration during the sheath tube contraction. The long residues 1-35 and 525-529 twist significantly more from the rest of the molecule making the latter turn as a single unit.

**Movie S3**. The atomic structure of the tail tube with one BplB subunit is colored in violet. The residues and their charges of the surrounding BplB molecules responsible for interacting with the colored subunit are shown. Red represents the negatively charged residue while blue represents the positively charged ones.

**Movie S4.** The atomic structure of the baseplate formed by a total of 10 unique gene products, each colored separately. The colors for each of the proteins are shown below.

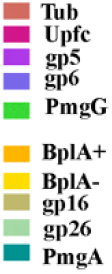

**Movie S5.** Interaction (H bonds) between the triplex molecule of the baseplate Hub form gp16 BplA+ and BplA-shown in red color.

**Movie S6.** A single sheath protofilament wrapped around the tail tube, both shown at atomic resolution, with the former undergoing a uniform upward contraction. The gap between the tail tube and sheath subunits increases as the protofilament compresses.

**Movie S7.** A single sheath protofilament wrapped around the tail tube undergoing a downward nonuniform cork-screw motion. The tail tube also follows a downward motion for penetrating the bacterial membrane for release of the phage genome for replication.

